# Ecogenomics reveals distinctive viral-bacterial communities in the surface microlayer of a natural surface slick

**DOI:** 10.1101/2023.02.24.528798

**Authors:** Janina Rahlff, Matthias Wietz, Helge-Ansgar Giebel, Oliver Bayfield, Emelie Nilsson, Kristofer Bergström, Kristopher Kieft, Karthik Anantharaman, Mariana Ribas-Ribas, Oliver Wurl, Matthias Hoetzinger, Alfred Antson, Karin Holmfeldt

## Abstract

Visible surface films, termed slicks, can extensively cover the sea surface, particularly in coastal regions. The sea-surface microlayer (SML), the upper 1-mm at the air-water interface in slicks (slick SML) harbors a distinctive bacterial community, but little is known about SML viruses. Using flow cytometry, metagenomics, and cultivation, we investigated viruses and the bacterial community from a brackish slick SML in comparison to non-slick SML as well as the seawater below (SSW). We conducted size-fractionated filtration of all samples to distinguish viral attachment to hosts and particles. The slick SML contained higher abundances of virus-like particles, prokaryotic cells, and dissolved organic carbon compared to non-slick SML and SSW. The community of 428 viral operational taxonomic units (vOTUs), 426 predicted as lytic, distinctly differed across all size fractions in the slick SML compared to non-slick SML and SSW. The distinctness was underlined by specific metabolic profiles of bacterial metagenome assembled genomes and isolates, which revealed prevalence of motility genes and diversity of CAZymes in the slick SML. Despite overall lower diversity, several vOTUs were enriched in slick SML over slick SSW. Nine vOTUs were only found in slick SML and six of them were targeted by slick SML-specific CRISPR spacers likely originating from Gammaproteobacteria. Moreover, isolation of three previously unknown lytic phages for *Alishewanella* sp. and *Pseudoalteromonas tunicata*, representing abundant and actively replicating slick SML bacteria, suggests that viral activity in slicks can contribute to biogeochemical cycling in coastal ecosystems.

## Introduction

The air-sea interface spans 70 % of Earth’s surface area and contains a diverse microbial community referred to as neuston [1], globally constituting 2×10^23^ cells [2]. In the uppermost 1-mm layer of the water column, termed the sea-surface microlayer (SML), the inhabiting organisms and viruses encounter highly dynamic conditions. The SML has been considered an “extreme” habitat influenced by high solar radiation, wind-wave interaction, accumulation of pollutants, sudden changes in temperature and salinity, contact to rainfall and depositions from the atmosphere, among other parameters [3–8]. While abundance, diversity and function of eukaryotes, bacteria, and archaea [9–12] have been studied in the SML, little is known about residing viruses (reviewed by Rahlff [13]). This is particularly true for the SML within natural surface slicks. Surface slicks form during low wind speeds and represent areas of accumulating surfactants, which by exceeding an unknown concentration threshold dampen small-scale capillary waves, resulting in a coherent surface film (Fig. 1a, [14,15]). Surface slicks are widely distributed with greater prevalence in coastal regions compared to the open ocean (covering on average 30 versus 11 % of surface area) but can occasionally cover the surface to up to 80 % in coastal waters (Romano 1996). Slicks are often enriched in cyanobacteria such as *Trichodesmium* [16–18], shown to decrease salinity and increase warming of the surface slick water [19]. Slicks also function as nurseries and dispersal agents for higher trophic levels [20,21], plus having an important function in the suppression of air-sea carbon dioxide fluxes [22,23]. Despite being little understood as microbial habitats to date (reviewed by Voskuhl and Rahlff [24]), slicks can accumulate and spread bacteria [25], and the bacterial community of slick SML remarkably differs from that of non-slick SML [26]. Based on 16S rRNA fingerprinting, Stolle, et al. [27] reported different particle-associated and free-living bacterial communities in the Baltic Sea during formation and disintegration of a surface slick, with strong community changes among free-living bacteria during slick disintegration. Other studies reported the presence of surfactant-producing bacteria like *Bacillus* spp. and *Pseudomonas* spp. in and under the SML within natural surface slicks [28–30].

**Fig. 1:**
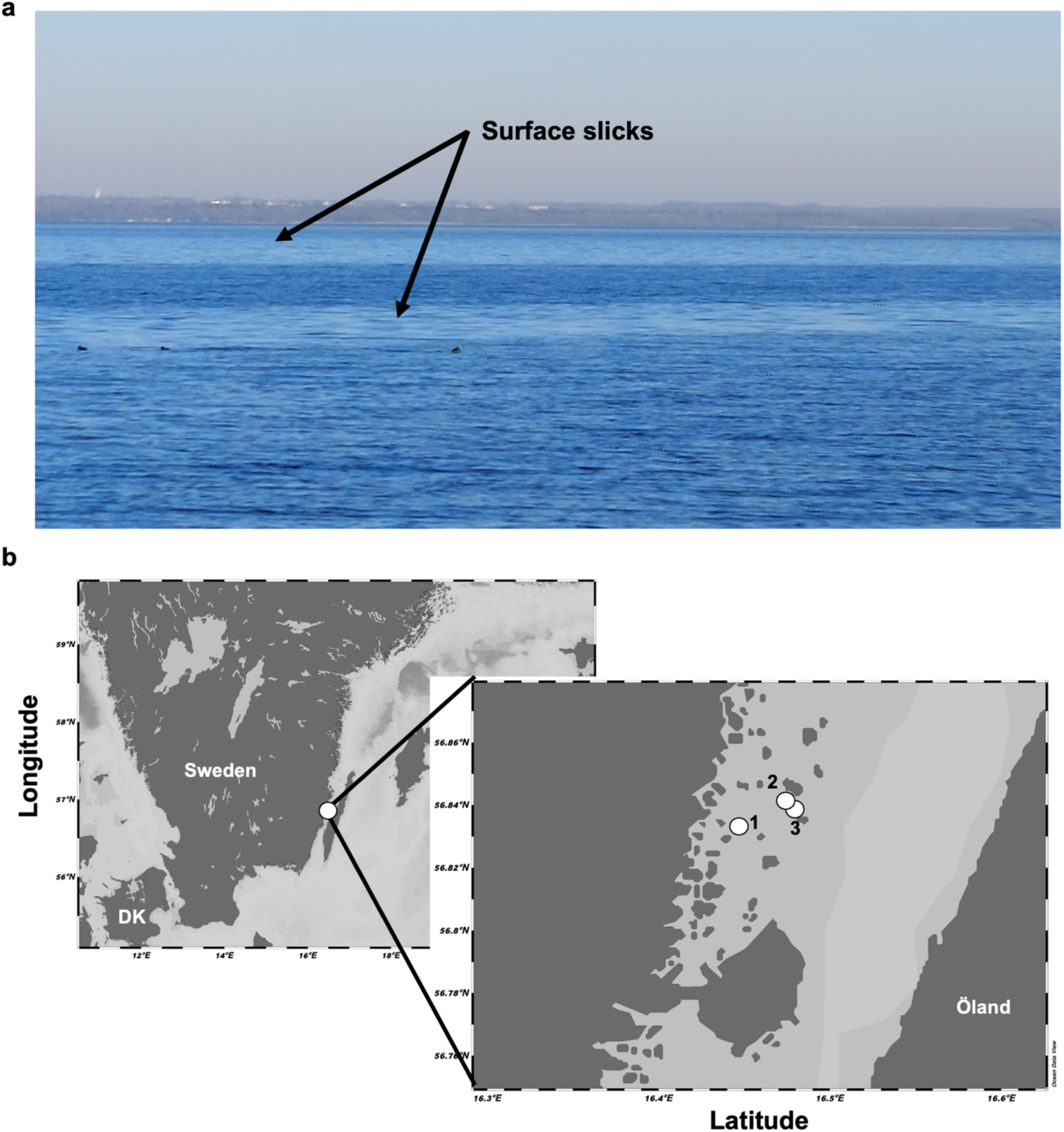
Representative example of surface slicks (none of the ones sampled), observed in the Kalmar Sound with Öland in the background **(a)**. Map illustrating slick sampling sites 1, 2, and 3 close to Ljungnäs (Rockneby, Sweden) in the Baltic Sea. Map was generated using Ocean Data View v.5.6.2 [116] **(b)**.

Investigations on viruses in non-slick SML reported increased virus-mediated mortality, increased virus-like particle abundance, and higher virus-host ratios compared to underlying water [31,32]. Work on Lake Baikal described autochthonous bacteriophage communities establishing in the microlayer [33], but such studies are lacking for marine systems. A pressing question in virioneuston research is whether viruses respond to harsh environmental conditions at the air-sea boundary by lysogeny, or if high host abundances favor lytic infections according to the “kill-the-winner” model, where viruses periodically decimate the most abundant hosts [34,35]. Currently there has been evidence for both, *i.e*. a predominant lytic viral activity in the SML [36] and the prevalence of prophages in the SML compared to underlying water [37]. In addition, viral abundance and diversity are unknown in surface slicks, and a comparison to non-slick SML is missing. Slicks often accumulate foams [26], and increased virus-like particle numbers have been shown for particle-enriched sea foams floating on the SML [32]. Slick SML is enriched with transparent exopolymer particles [26], absorbing viruses [38], but to which extent viruses, particularly bacteriophages, are associated with particles within slicks is unknown. The SML is a net heterotrophic system [39], as primary producers suffer from photoinhibition [40]. During the “viral shunt” [41] when viruses lyse their hosts and dissolved organic matter enters the water column, heterotrophy might be further supported in the SML. By contributing to release of surfactants and organic matter, the virioneuston could effectively contribute to the biological carbon pump and biogeochemical cycling.

In this work, we combined metagenomics and cultivation to reveal viral-bacterial dynamics in the SML of a surface slick from the coastal Baltic Sea compared to non-slick SML and underlying surface water. We accounted for the role of virus-host and virus-particle associations via size-fractionated filtration of water samples. Furthermore, by studying patterns in the clustered-regularly interspaced short palindromic repeats (CRISPR)-Cas systems – the adaptive immune systems of bacteria – we revealed past infection histories in the SML microbiome.

## Materials and methods

### Sampling

Water samples inside and outside a surface slick from site #1 – #3 were collected from a small boat in Kalmar Sound, at Ljungnäs near Rockneby, Sweden on the 31^st^ of May 2021 (Fig. 1b and Table S1). The SML from slick and non-slick areas was collected using the glass plate method [42]. Reference water (subsurface water = SSW) was collected from ~70 cm depth below the slick and non-slick area with a Hydro-Bios 1-l Ruttner water sampler (Swedaq, Höör, Sweden). Wind speed was measured with a hand-held anemometer model MS6252A (Mastech Group, Brea, CA, USA). Site #1 was sampled for metagenomics, dissolved organic carbon (DOC), surfactants, and bacterial isolation, while sites #1 – #3 were sampled for flow cytometry and phage isolation (Table S1). All water samples (600 – 700 and 2100 ml of SML and SSW, respectively) were sequentially filtered through 5 μm pore size to obtain the particle-associated (PA) fraction and collecting the filtrate on 0.2 μm pore size to obtain free-living (FL) fraction using polycarbonate filters (Nucleopore Track-Edged Membrane, Whatman, Maidstone, UK). The PA fraction was unfiltered and assumed to contain > 5 μm microbes including phytoplankton, protists, virions attached to particles or hosts as well as intracellular viruses, compared to microbes between 5 and 0.2 μm size including any host-associated viruses in the FL fraction. The flow-through of the 0.2 μm filtration was chemically flocculated [43] using a higher iron-III-chloride concentration (10 g l^-1^) than in the original protocol as recently suggested for freshwater [44], and filtered onto 1 μm polycarbonate membrane filters (142 mm diameter, Whatman/GE Healthcare, Uppsala, Sweden) representing the viral fraction. All filter membranes were stored at −80 °C until DNA extraction.

### Surfactant and DOC analysis

Surfactant concentration was measured by the voltammetry 747 VA Stand (Metrohm, Herisau, Switzerland) with a hanging mercury drop electrode. Surfactants accumulate at the mercury drop at a potential of −0.6 V /versus an Ag/AgCl reference electrode. Surfactants were quantified in 10 ml of unfiltered samples with the standard addition technique with details given in [45]. For DOC, duplicates of 30 ml sample water and a MilliQ control were gravity-filtered onto precombusted (475 °C, 3 h) GF/C glass fiber filters (nominal pore size ~1.2 μm), acidified (200 μl 2 M HCl), and stored in precombusted glass vials (475 °C, 3 h) with acid-washed lids at 4 °C until analysis as described previously [46].

### Cell count and virus-like particle measurements

Unfiltered slick SML, non-slick SML, slick SSW and non-slick SSW from sites #1 – #3 (Fig. 1b) were fixed with 25 % glutardialdehyde (0.5 % final concentration, Sigma-Aldrich/Merck Life Science AB, Solna, Sweden) and stored at −80 °C. Unfortunately, the samples experienced an extra freeze-thaw cycle during shipment, but we are confident that comparisons between samples treated the same are reasonable. Prokaryotic cells and virus-like particles (VLPs) from site #1 – #3 were measured on a flow cytometer (BD Accuri C6, BD Biosciences, Franklin Lakes, NJ, USA) by using protocols from [47] and [48], respectively. Enrichment factors (EF) were calculated for flow cytometry data and metagenome coverages of viral OTUs (see below) as a ratio of a factor in the SML divided by the SSW counterpart, with EF > 1 and EF < 1 indicating enrichment and depletion in the SML, respectively.

### Isolation of bacteria

Bacteria were isolated from site #1 by plating water on Zobell Agar (1 g yeast extract (BD), 5 g bacto-peptone (BD), 15 g bacto agar (BD), 800 ml Baltic Sea water, 200 ml Milli-Q water). Plates were incubated at room temperature. Selected bacteria were pure-cultured from single colonies thrice before they were inoculated in Zobell medium (1 g yeast extract (BD), 5 g bacto-peptone (BD), 800 ml Baltic Sea water, 200 ml Milli-Q water) over night, and stored as glycerol stocks (600 μl 50% glycerol (Sigma) and 900 μl bacterial culture) at −80 °C.

### Phage isolation and plaque assay

Water from all sampling sites was filtered through a 0.2 μm PES syringe filter and the flow-through collected for phage isolation using plaque assay following Nilsson, et al. [49]. Briefly, 500 μl of the water sample was mixed with 3.5 ml top agar (450 mM NaCl (Sigma), 50 mM MgSO_4_ x 7 H_2_O (Sigma), 50 mM Trizma base (Sigma), 5 g l^-1^ low-melting agarose (Thermo Scientific, Waltham, MA, USA)) and 300 μl overnight bacterial culture. Plates were incubated on the bench overnight, and plaque-forming units were monitored over 48 h. Selected phage plaques were picked from plates using a sterile 100 μl pipet tip and stored in MSM buffer (= top-agar without low-melting agarose) at 4 °C. Phages were purified by replating thrice before two full lysed plates per viruses was harvested with 5 ml MSM buffer. The phage-MSM mixture was centrifuged at 4500 rpm for 20 min., and the supernatant was filtered through a 0.2 μm syringe filter and stored at 4 °C. The phages were stored both as free phages at 4 °C and in infected hosts at −80 °C. For infected hosts, the 400 μl freshly harvested phage stock were mixed with 1.2 ml overnight bacterial culture for 15 min. before being mixed with glycerol and frozen as described above.

### Transmission Electron Microscopy (TEM) imaging

TEM was conducted using high titer phage lysate and negative staining as in [50]. Briefly, phages were loaded on pre-discharged copper grids (200 Mesh Cu. Agar Scientific Ltd., Stansted, UK), stained with 2 % w/v uranyl acetate (Agar Scientific Ltd), and imaged using an FEI Tecnai 12 G2 BioTWIN microscope. Phage head and tail diameter were measured with ImageJ v.1.53 [51] according to [52].

### DNA extraction and sequencing of bacterial and phage isolates and metagenomes

DNA from 1 ml of harvested phage stock was extracted using Wizard PCR DNA Purification Resin and Minicolumns (both Promega, Madison, WI, USA) as described previously [49]. Bacterial genomic DNA was extracted using the E.Z.N.A Tissue DNA kit (Omega Bio-tek, Norcross, GA, USA) according to the manufacturer’s protocol. DNA for metagenomes was extracted from 5 and 0.2 μm filters (47 mm diameter) using the DNAeasy Power Soil Pro kit (Qiagen, Sweden). DNA from viral fraction (142 mm diameter membranes) was extracted using the DNAeasy PowerMax Soil kit (Qiagen, Sweden) and a subsequent ethanol precipitation step for concentrating DNA. DNA concentrations were measured on a Nanodrop 2000 spectrophotometer (Thermo Scientific) and Qubit^®^ 2.0 Fluorometer (Invitrogen/ Life Technologies Corporation, Carlsbad, CA; USA). Sequencing was conducted by SciLifeLab (Solna, Sweden) using Illumina DNA PCR-free library preparation and an Illumina NovaSeq6000 sequencer using a NovaSeq S4 flowcell (2 x 150 bp). One bacterial (SMS8) and one viral (vB_PtuP_Slicky01) genome were sequenced at Eurofins Genomics using the INVIEW Resequencing bacteria (eurofinsgenomics.eu) product and the same sequencer as above. All reads went through adapter trimming and quality control using bbduk as part of BBTools [53] and Sickle [54]. Genomes from isolates were assembled using MEGAHIT v.1.2.9 for phages [55] and SPAdes v.3.15.3 with option --isolate for bacteria [56]. All reads went through adapter trimming and quality control using bbduk as part of BBTools [53] and Sickle [54]. Genomes from isolates were assembled using MEGAHIT v.1.2.9 for phages [55] and SPAdes v.3.15.3 with option --isolate for bacteria [56]. Quality checks and taxonomy assignments were performed as for metagenome-assembled genomes (MAGs), see below. Several functional categories in genomes of SML bacterial isolates were annotated (Table S2a-g). KEGG annotations were done using KAAS [58], and pathways reconstructed using KEGG Mapper [59]. CAZymes were predicted using dbCAN2 [60] only considering hits with > 60 % query coverage and e-value < 1e-15. Analyses were done and visualized in R v.4.2.2 using the package tidyverse [61]. Biosynthetic gene clusters were predicted using antiSMASH v.6.1.1 [62]. Genes involved in surfactant biosynthesis (NCBI accessions AAD04757.1, AEW31038.1, NP_252169.1, PBL99399.1, BAG28347.1, AAB35246.1) were searched using Custom-BLAST in Geneious Prime [63]. For comparison, the lichenysin gene cluster was obtained from https://mibig.secondarymetabolites.org [64].

### Binning and functional analysis of bacterial genomes

Taxonomic profiling of bacteria was conducted using mOTUs v.3.0.2. using trimmed reads [65]. Resulting read counts were read-sum normalized, and Shannon-Wiener index and relative abundance for beta-diversity (also for the viral clusters (VC), see below) were investigated using phyloseq package [66] in the R programming environment [67]. Binning of MAGs was performed using CONCOCT v.1.1.0 [68] and MetaBAT v.2.12.1 [69] on MetaSPAdes [56] v.3.15.3 assemblies previously filtered to a minimum length of 1000 bps. A non-redundant set of bins was created with DAS_Tool v.1.1.3 [70] with default score threshold and followed by manual refinement in uBin v.0.9.14 [71] using information on GC content, coverage and taxonomy. MAGs underwent quality checks in CheckM v.1.1.3 [72], followed by taxonomic classification with the classify_wf option in GTDB-Tk v.1.7.0. and database version r202 [73]. MAGs were used for further analysis if they reached estimated completeness and contamination scores of ≥70 % and ≤10 % in either uBin or CheckM. Mapping to MAGs and isolate genomes was performed with Bowtie v.2.3.5.1 [74] using the --reorder flag. Mismatch filtering with 2% error rate (-mm 3) was conducted within iRep v.1.1.0, which was used to estimate *in situ* replication rates at default thresholds [75]. Average nucleotide identity (ANI) comparison was carried out using FastANI v.1.33 [76] and visualized in ANIclustermap (https://github.com/moshi4/ANIclustermap). KEGG annotations derived from predictions with DRAM v.1.2.4 [57] were compared for significant differences between MAG groups using ALDEx2 [77]. The number of hits were normalized by the number of genes per MAG. CAZymes were predicted as mentioned above for bacterial isolates.

### Metagenomic analyses of dsDNA viruses

Trimmed reads were assembled twice using MetaSPAdes v3.15.3 [56] and the Metaviral SPAdes [78] option. For viral analysis, assemblies were combined, filtered to keep scaffolds of minimum 1 kb, and viruses were identified using VIBRANT v.1.2.1 [79] by adding the --virome option for the viral fractions, and VirSorter v.2 with --include-groups “dsDNAphage,ssDNA” and default score [80]. The output was combined and filtered to 10 kb sequence length. Only viruses with attributes “medium quality”, “high quality” or “complete” determined using CheckV v.0.8.1 were used for downstream analyses. VIRIDIC v1.1 [81] was run for genus and species clustering, and only one representative of a viral species cluster (preferably a circular or the longest scaffold of the cluster) was used for further analysis. This workflow resulted in 428 representative viral scaffolds (further referred to as viral operational taxonomic units = vOTUs) as the final output. Viral relative abundance (depth of coverage) and breadth of coverage for vOTUs and phage isolates (see below) was calculated with scripts and settings as done previously [82] after read-mapping with Bowtie2 and following conventions of [83]. Viral genes in vOTUs and phage isolates were predicted using Prodigal v.2.6.3 in meta mode [84] and functionally annotated using DRAM-v v. 1.2.4 [57]. Synteny of selected vOTUs was visualized using clinker v.0.0.25 [85]. Clustering of vOTUs with reference database phages (release July 2022, from https://github.com/RyanCook94/inphared) [86] was performed using vConTACT2 v.0.9.19 [87], VC information was compiled using graphanalyzer v.1.5.1. (https://github.com/lazzarigioele/graphanalyzer), and the network visualized using Cytoscape v.3.9 [88]. Auxiliary metabolic genes (AMGs) on vOTUs were detected using annoVIBRANT (https://github.com/AnantharamanLab/annoVIBRANT), using modified scripts of VIBRANT v.1.2.1 [79] to report AMGs of vOTUs. For this analysis, a vOTU carrying an AMG was counted towards a sample, if it was present based on read mapping conventions [83], and the sum of coverage of phages carrying that AMG was calculated. Comparisons of enrichments of a vOTU inside and outside the slick was done by calculating coverage ratios, i.e., the coverage of a virus in slick SML divided by the coverage in slick SSW and the same procedure for the non-slick vOTUs. EF was calculated for coverage of VCs as explained above.

### CRISPR analysis and virus-host matches

Viral OTUs were matched to a set of MAGs previously dereplicated with dRep v.3.4.0 under default parameters [89] using VirHostMatcher v.1.0.0 [90] and by applying a d2* dissimilarity threshold of 0.3. Prophages in dereplicated MAGs and isolate genomes were detected with VIBRANT. CRISPRcasFinder v.4.2.20 [91] was used to detect CRISPR arrays in MetaSPAdes assemblies (>1 kb), MAGs and genomes of bacterial isolates. CRISPR direct repeat (DR) sequences from assemblies were extracted from evidence level 4 CRISPR systems (Table S3a). DR sequences were blasted against vOTUs, and DRs with a hit at 100% similarity were deleted to avoid extraction of false-positive spacers from vOTUs. Remaining DRs (Table S3b) were fed into the Metagenomic CRISPR Reference-Aided Search Tool (MetaCRAST) [92] to extract spacer sequences from raw reads using settings -d 3 -l 60 -c 0.99 -a 0.99 -r. Spacers were subsequently filtered for homopolymers and length (20 – 60 bp), only considering CRISPR spacer to viral protospacers matches with 100 % similarity (very strict filtering due to high amount of matches). Spacer-protospacer matches between MAGs, bacterial isolates, and viruses were filtered at 80 % similarity.

## Results

### Surfactants, DOC, VLPs and prokaryotic cells are enriched in the slick SML

Slick SML was enriched in dissolved organic carbon (DOC, 7.69 and 7.83 mg l^-1^) in comparison to non-slick SML (5.31 and 5.40 mg l^-1^) and both SSW samples (5.08 and 5.09 mg l^-1^) (Fig. 2a). The same pattern was seen for surfactants in slick SML (mean = 1219.3 μg Triton-X-100 equivalent (Teq) l^-1^) compared to the other samples (334.3 – 412.1 Teq l^-1^, Fig. 2b). Prokaryotes and VLPs from three individual surface slicks showed highest counts in the slick SML compared to non-slick SML and SSW (Fig. 2c). Prokaryotic abundance in slick SML was 2.7 x 10^6^ ± 1.3 x 10^6^ cells ml^-1^ compared to 5.9 x 10^5^ ± 3.5 x 10^4^, 5.8 x 10^5^ ± 9.5 x 10^4^, and 5.8 x 10^5^ ± 3.7 x 10^4^ cells ml^-1^ in the non-slick SML and SSW samples, respectively. Abundances of VLPs in slick SML was 1.2 x 10^8^ ± 5.4 x 10^7^ compared to 1.4 x 10^7^ ± 9.5 x 10^6^, 2.0 x 10^7^ ± 9.5 x 10^6^, and 1.4 x 10^7^ ± 5.3 x 10^6^ VLPs ml^-1^ in the other samples (Fig. 2c). Virus-host ratios were highest in slick SML (44.6 ± 8.0), followed by slick SSW (34.5 ± 14.0), non-slick SML (23.4 ± 3.0) and non-slick SSW (23.3 ± 7.5). Mean EFs were 6.0 ± 1.0 and 1.1 ± 0.5 for VLPs in slick SML and non-slick SML, respectively, and 4.5 ± 1.4 and 1.0 ± 0.1 for prokaryotic cells for slick SML and non-slick SML, respectively.

**Fig. 2:**
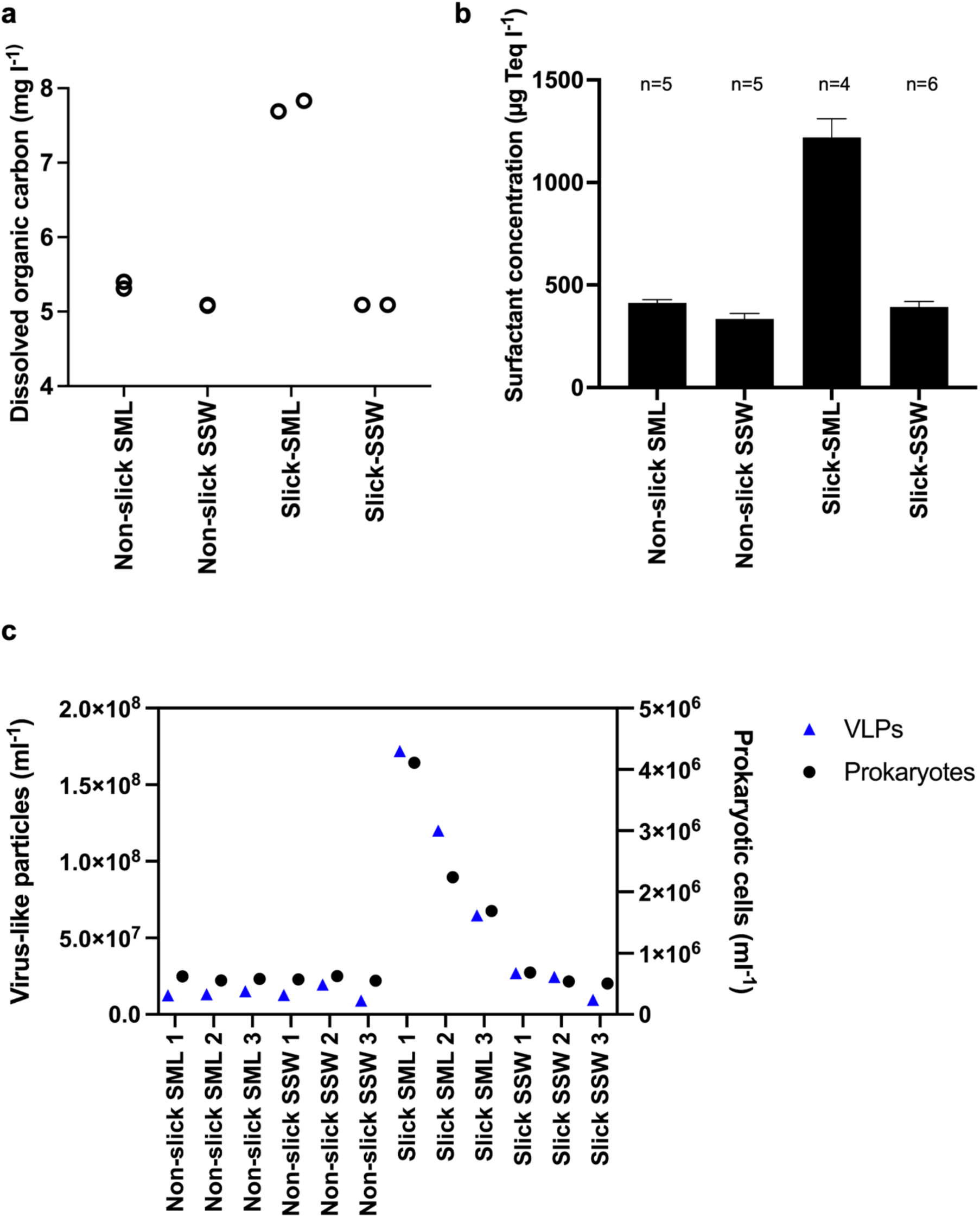
Dissolved organic carbon (DOC) measured in technical duplicates **(a)**, mean +/- standard deviation for concentration of surfactants (n=number of technical replicates) (**b)**, and counts of virus-like particles (VLP) and prokaryotic cells in slick SML, non-slick SML, slick SSW and non-slick SSW **(c)**. DOC and surfactants were measured from sampling site #1 only. SML = sea-surface microlayer, SSW = subsurface water (~ 70 cm depth)

### Higher bacterial diversity in slick SML with Gammaproteobacteria as dominant class

α-diversity was highest in the PA bacterial fraction (>5 μm) within the slick SML, illustrated by maximum Shannon-Wiener diversity index 3.7 compared to non-slick SML (3.0), slick SSW (3.2) and non-slick SSW (2.9). A similar, but weaker, trend was observed for the FL bacterial fraction (5 – 0.2 μm) with a Shannon-Wiener index of 3.1, 2.9, 2.9 and 2.8 for slick SML, non-slick SML, slick SSW, and non-slick SSW, respectively (Fig. 3a). Especially Gammaproteobacteria showed higher relative abundance in the slick SML in both the PA (38.5 %) and FL (48.8 %) fraction compared to other samples (7.5 – 15.1 %, Fig. 3b&c). Among the Gammaproteobacteria, most abundant families in the slick SML FL and PA fraction were *Pseudoalteromonadaceae*, mostly *Pseudoalteromonas tunicata* (22.1 % compared to 0.01 % in non-slick SML) and *Chromatiaceae* (17.8 % compared to 1.2 % in non-slick SML, Fig. 3c, Supplementary Results), respectively. In the slick SML PA fraction, other abundant bacteria were *Polaribacter* spp. (8.3 %, Fig. S1), *Nodularia spumigena* (4.2 %), *P. tunicata* (3.8 %), *Pseudomonas fluorescens* (2.9 %), and *Shewanella baltica* (2.5 %). The slick SML FL fraction featured unclassified Verrucomicrobia (10.7 %) and *Marinomonas* (8.4 %) species. On the other hand, unclassified *Porticoccaceae* were less abundant in the slick SML (3.6 %) compared to non-slick SSW (7.3 %) in the FL fraction.

**Fig. 3:**
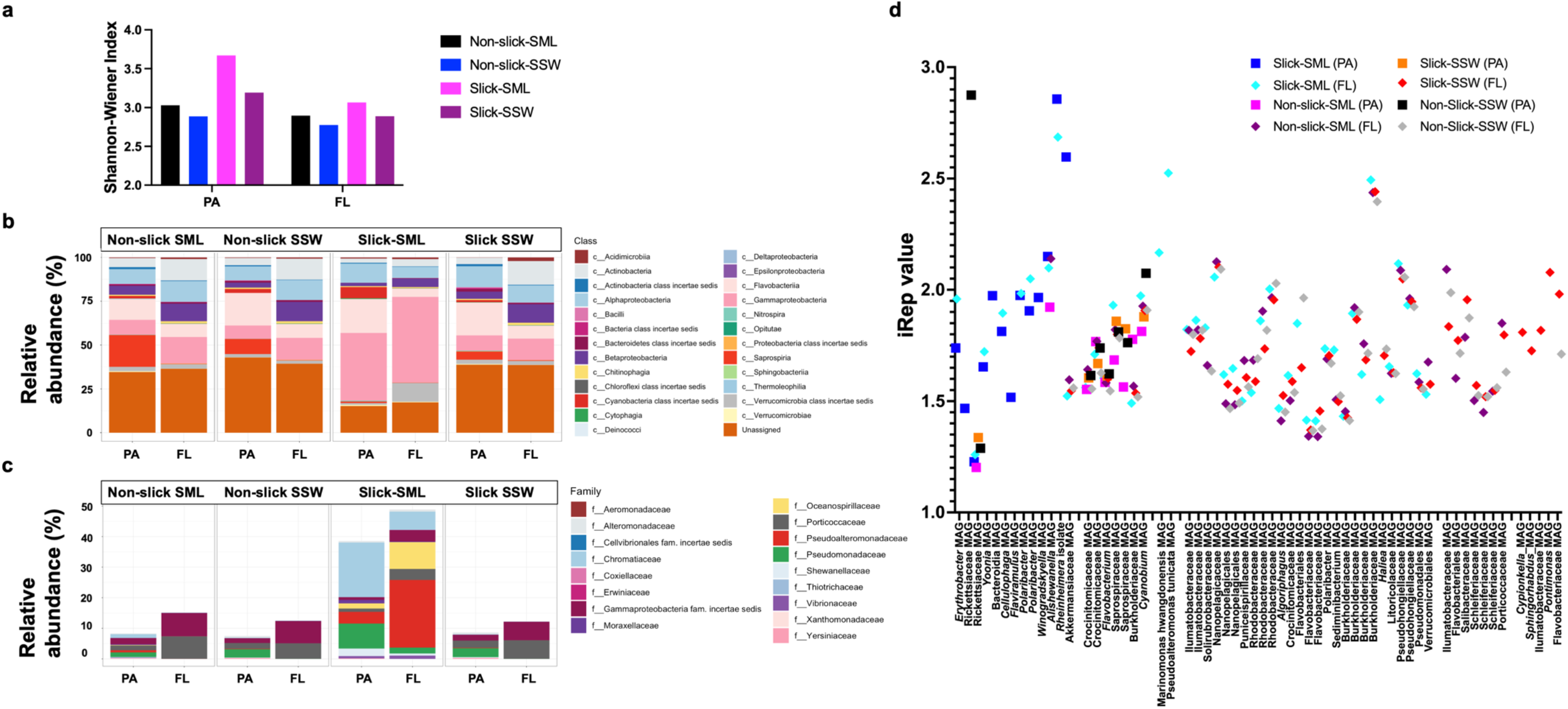
Diversity and indices of replication (iRep) for slick-associated bacteria. Shannon-Wiener Index for PA and FL bacterial fractions from slick versus non-slick samples **(a)**, relative abundance of bacterial classes among PA and FL fractions **(b)**, and families of Gammaproteobacteria in close-up **(c)**. In situ replication rates (based on iRep) for bacterial metagenome-assembled genomes **(d)**. FL = free-living fraction (5 – 0.2 μm pore size filtered), PA = particle-associated fraction (> 5 μm filtered), SML = sea-surface microlayer, SSW = subsurface water (~ 70 cm depth).

### Abundant bacteria with predicted activity reside in the particle-associated fraction of the slick SML

We recovered 316 MAGs and seven gammaproteobacterial isolates from slick SML samples and performed functional predictions to assess putative ecological traits. The seven strains were sequenced: *P. tunicata* SMS2, *Alishewanella* sp. SMS8, SMS9, as well as *Rheinheimera baltica* SMS3, SMS4, SMS11, SMS12 (Table S2a). The 316 MAGs covered eight bacterial phyla (Table S4a). Four MAGs carried prophages: MAG_221 (*Flavobacteriaceae*), MAG_166 (*Rickettsiaceae*), MAG_147 (*Cypionkella* sp.), and MAG_137 (*Cyanobium* sp.), but among these four, only MAG_166 showed higher coverage (101 x) in slick SML compared to other samples (Table S4a&S5). Analysis of *in situ* replication rates represented by the Index of Replication (iRep) suggested that several MAGs and the genome of SMS3 (representative genome of *R. baltica*) formed a distinct group of actively replicating bacteria in the slick SML PA fraction (Fig. 3d). An iRep > 2 in the PA fraction was found for MAGs of *Alishewanella, Akkermansiaceae*, the *R. baltica* SMS3 (Fig. 3d, Fig. S2) matching their high relative abundance in the PA fraction (Table S5). Analysis of iRep for MAGs of *Marinomonas hwangdonensis* (2.2) and *P. tunicata* (2.5) suggested that these bacteria replicated in the FL fraction of the slick SML. MAG_01 as well as *Alishewanella* isolates SMS8 and SMS9 formed a joint ANI cluster (ANI ≥ 99.3 %) (Fig. S3), and SMS8 and SMS9 are probably new species and assigned to *Alishewanella* sp. in GTDB-Tk classify workflow (Fig. S4). SMS8 carried a prophage likely with siphovirus morphology and a genome length of 50 kb (Table S2a, Fig. S5, Supplementary Results).

### Functional characterization of MAGs and isolates

MAGs and isolates abundant in slick SML (Table S5) were functionally analyzed regarding their diversity using KEGG modules and CAZymes, informing about central metabolic capacities and carbohydrate degradation, respectively. To identify slick- and SSW-specific features, four MAGs predominant in slick at high abundance (designated Slick_highAb), eight MAGs predominant in slick at low abundance (designated Slick_lowAb), and five MAGs predominant in the SSW (designated SSW, Table S4b) were analyzed. Approximately 250 KEGG-ids were differentially abundant between the three groups (Table S4c). For instance, flagellum genes were only found in Slick_highAb MAGs, and several genes involved in regulation, repair and biosynthesis showed contrasting patterns between groups (Fig. 4a). SSW MAGs encoded more CAZymes (Fig. S6, Table S4d), with significantly higher fractions of glycoside hydrolase, polysaccharide lyase, and glycosyltransferase genes (Wilcoxon test, *p* < 0.05). However, the diversity of CAZyme families was higher in SML-MAGs, with complementing CAZyme profiles between Slick_lowAb and Slick_highAb MAGs (Fig. 4b). Both groups of SML-MAGs encoded diverse polysaccharide lyase families, whereas the SSW group only encoded alginate-targeting PL6, PL7 and PL17 but in higher copy numbers. However, sizes of gene pools differed, with 9000 genes in SSW-MAGs compared to ~60000 in SML-MAGs, possibly influencing these patterns.

**Fig. 4:**
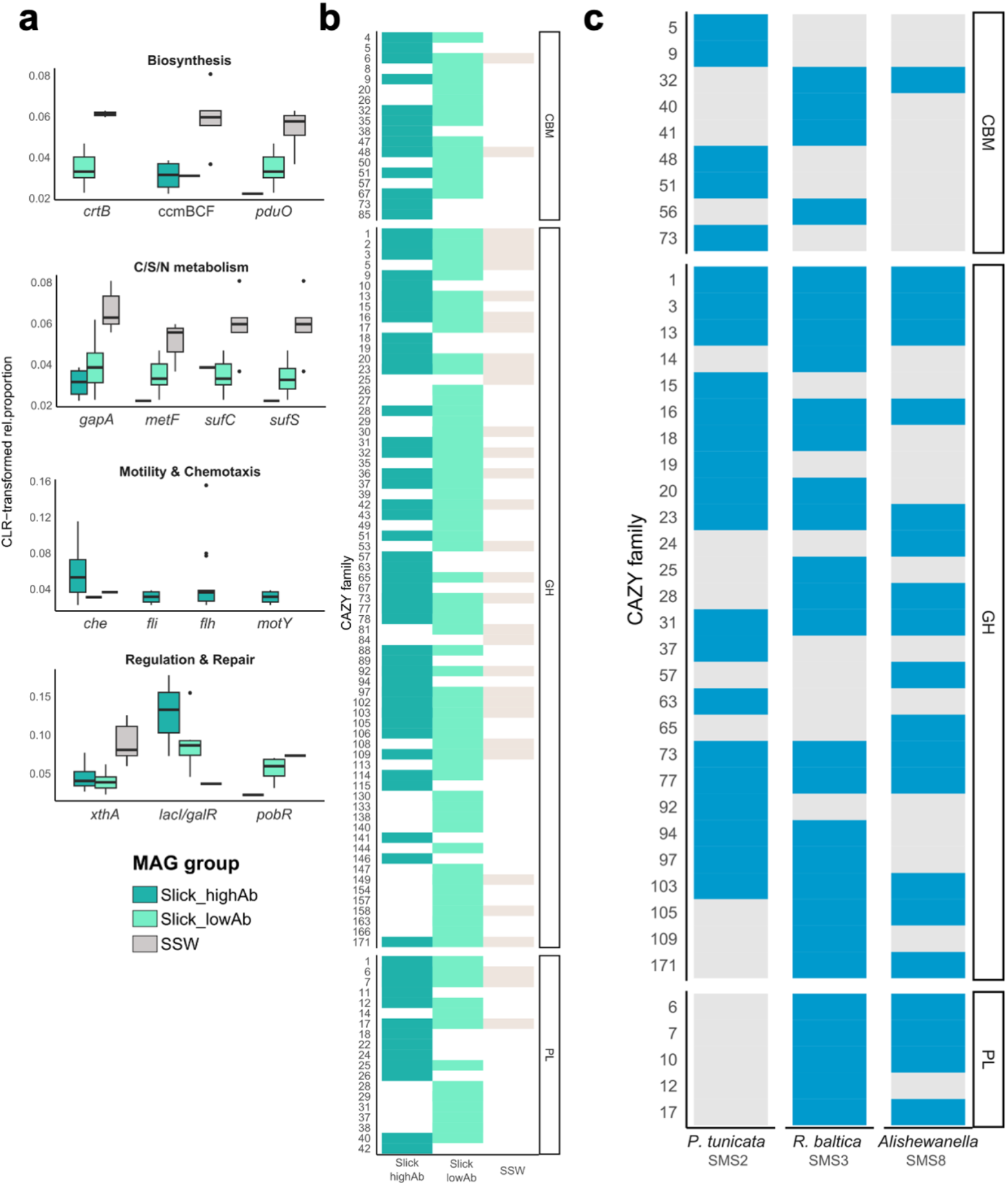
Relative fraction of genes from specific functional pathways with differential abundance between MAG groups, identified using Aldex2 and displayed a CLR-transformed relative gene abundances. Several genes from functionally related categories (e.g. che chemotaxis, fli/flh flagellum genes) were combined, showing the average CLR value. lacI / galR: LacI family transcriptional regulator; xthA: exodeoxyribonuclease; motY: sodium-type flagellar protein; pobR: AraC family transcriptional regulator; sufS: cysteine desulfurase / selenocysteine lyase; sufC: Fe-S cluster assembly ATP-binding protein; metF: methylenetetrahydrofolate reductase; gapA: glyceraldehyde 3-phosphate dehydrogenase; pduO: cob(I)alamin adenosyltransferase; crtB: 15–cis–phytoene synthase **(a).** Diversity of CAZyme families between MAG groups (see Table S2G for details) **(b)**. CAZyme profiles of slick SML isolates, showing presence/absence of carbohydrate-binding module (CBM), glycoside hydrolase (GH), and polysaccharide lyase (PL) gene families. The four R. baltica isolates featured identical CAZyme diversity; therefore, only SMS3 is shown as representative. Due to lower completeness of SMS9, only SMS8 is shown for Alishewanella sp. **(c)**.

To corroborate the distinctness of SML microbiomes, we analyzed the genomes of the seven strains from slick SML, for which corresponding MAGs have a slick-specific high abundance and iRep compared to SSW (Table S2a, S4b, S6a). Due to low completeness of SMS9, we focused on SMS8. All isolates shared multiple gene clusters mediating chemotaxis and motility (e.g, mcr/che, mot, fli; Table S2d). *P. tunicata* SMS2 and *R. baltica* SMS3, SMS4, SMS11, and SMS12 additionally harbored a type VI secretion system involved in biofilm formation. All *R. baltica* strains featured highly similar CAZyme and KEGG profiles, with only 58 of 7850 predicted KEGG-ids not shared. We found genes encoding glycoside hydrolases GH103 and GH171 in *R. baltica* (Fig. 4b) and the same genes for homoserine lactone, mediators of quorum sensing, in all *R. baltica* and another two in *Alishewanella* sp. SMS8 (Table S2f). SMS2 encodes a distinct CAZyme repertoire compared to other strains (Fig. 4c), indicating that SML strains specialize on different carbon sources. Most notably, SMS2 encodes no polysaccharide lyases compared to several clusters targeting alginate and pectin in the other strains. Instead, the unique presence of GH19 plus carbohydrate-binding module families CBM5 and CBM73 might enable chitin degradation; plus mannan, amylase and pullulan activities through GH92 and GH13 genes. Overall CAZyme profiles of SMS2 and SMS3 sometimes complemented each other (Fig. 4c), suggesting that the strains occupy different trophic niches. SMS2 encoded genes for violacein and prodigiosin biosynthesis, likely explaining its purple-blue phenotype (Table S2a). Checking for surfactant-related genes revealed several homologs of lichenysin and surfactin synthetase (30 % amino acid identity to genes in the characterized cluster of *Bacillus licheniformis*) in strains SMS2, SMS3, and SMS4 (Table S2g).

### Viruses establish a distinctive community in the slick SML

Of 428 vOTUs > 10 kb length (dsDNA viruses) dereplicated at the species level, 16, 45, and 367 were complete, high, or medium quality, respectively. Only two vOTUs were determined to be proviruses according to CheckV (Table S6). Different vOTUs were assembled in each sample type and size-fraction but based on read mapping, most vOTUs and VCs were shared between all four sample types (Fig. S7 & S8). Certain vOTUs were unique for the slick SML, while others were found in all sample types except slick SML (Fig. S7) agreeing with a correlation matrix showing that the slick SML vOTU community was most distinct from the other samples (Fig. S9), and virus coverage was highest in the viral fractions (Fig. S10). Contrasting the bacterial α-diversity, vOTUs were more diverse in the FL and viral fraction (Shannon-Wiener index: range 3.4 – 4.6) compared to PA (range 2.5 – 3.8) fractions, but among each filtered fraction always lowest in slick SML compared to the other samples (PA = 2.6, FL = 3.7, viral = 3.5, Fig. 5a). Certain viral clusters (VCs) showed markedly higher relative abundance in slick SML such as VC_988_0 with relative abundance of ~43 % (slick SML) compared to ~8 % (non-slick SML) (Fig. 5b). According to the vConTACT2 analysis, VC_988_0 shares protein clusters with Flavobacteria phages (Fig. 5c cluster 1). In the PA fraction, vOTUs of VC_1425_0 carrying a reverse transcriptase gene showed a relative abundance of 51.5 % in slick SML compared to 3.3 % in non-slick SML. Nine vOTUs were exclusively detected in the slick SML. Among those, the overlap cluster VC_580/VC_601, sharing genomic similarities with various VCs containing known *Shewanella* phages, contained two vOTUs (34.1 kb and 40.7 kb length) that were solely detected in slick SML PA fraction (Fig. S7g).

**Fig. 5:**
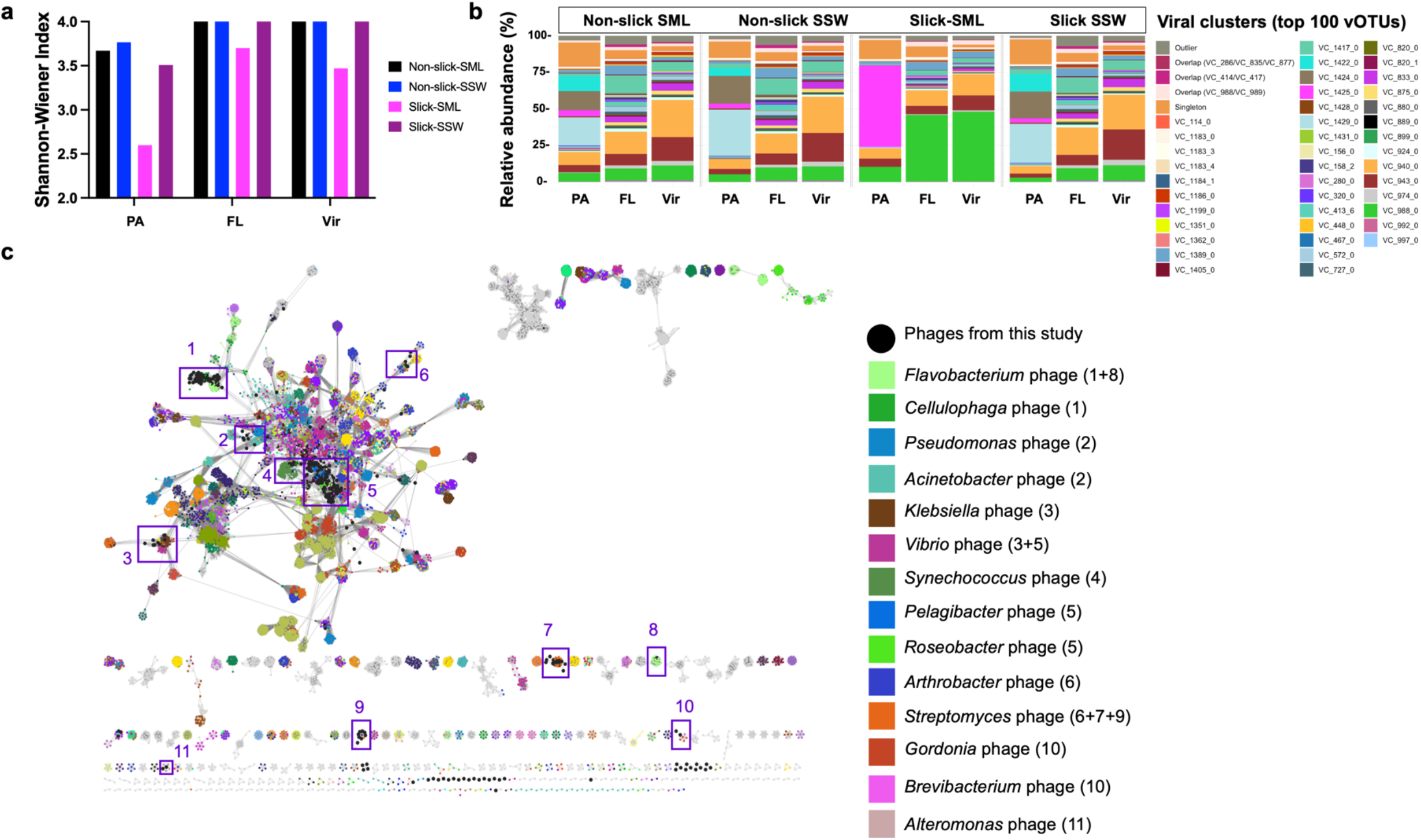
Viral diversity and clustering. Shannon-Wiener Index for vOTUs for the four different sample types **(a)**. Relative abundance of viral clusters (VC) including the top 100 abundant vOTUs show an increase in relative abundance of certain VC in slick SML (VC_1425_0 & VC_988_0), while other VC decreased in relative abundance (VC1424_0 or VC_572_0) compared to reference samples. Further information about VCs and closest associated viruses is given in Table S6. Outliers and singletons refer to unclustered, presumably unknown viruses **(b)**. Many vOTUs (nodes) from this study clustered with known phages such as Flavobacterium, Pelagibacter, or Synechococcus phages based on shared protein clusters (interactions with known phages indicated by eleven purple frames) with a virus reference database from July 2022. Several vOTUs clustered only with other vOTUs from this study indicating unknown viruses **(c)**. FL=free-living fraction (5 – 0.2 μm pore size filtered), PA=particle-associated fraction (>5μm filtered), SML=sea-surface microlayer, SSW=subsurface water (~70 cm depth), Vir = viral fraction (< 0.2 μm)

Less VCs became enriched (EF > 1) in the PA and FL fraction of the slick SML compared to non-slick SML (Fig. 6a), in agreement with the α-diversity (Fig. 3a). However, the few vOTUs that were enriched in the slick SML often reached very high EFs (> 6), *e.g*., subclusters of VC_988_0, VC_975_0 (resembling *Pelagibacter* phage HTVC028P), VC_1075_0 (resembling *Gordonia* phage GMA6), VC_1182, VC_1425_0, as well as several singletons and outliers, presumably representing previously unknown viruses (Table S6 & S7). Likely due to their higher abundance, viruses contributed to prevalence of more viral AMGs in the slick SML (Fig. 6b, Table S8 a&b, Fig. S11), namely genes related to amino acid metabolism (mainly arginine, proline, alanine, aspartate and glutamate metabolism), carbohydrate metabolism (amino sugar, nucleotide sugar, fructose and mannose metabolism) or to cofactors and vitamins (e.g. folate biosynthesis). The abundant 106.6 kb and 57.4 kb vOTUs from the slick SML, both unclustered in vConTACT2, contained the gene *folA* (dihydrofolate reductase (KEGG enzyme EC: 1.5.1.3), which has an essential role in DNA synthesis.

**Fig. 6:**
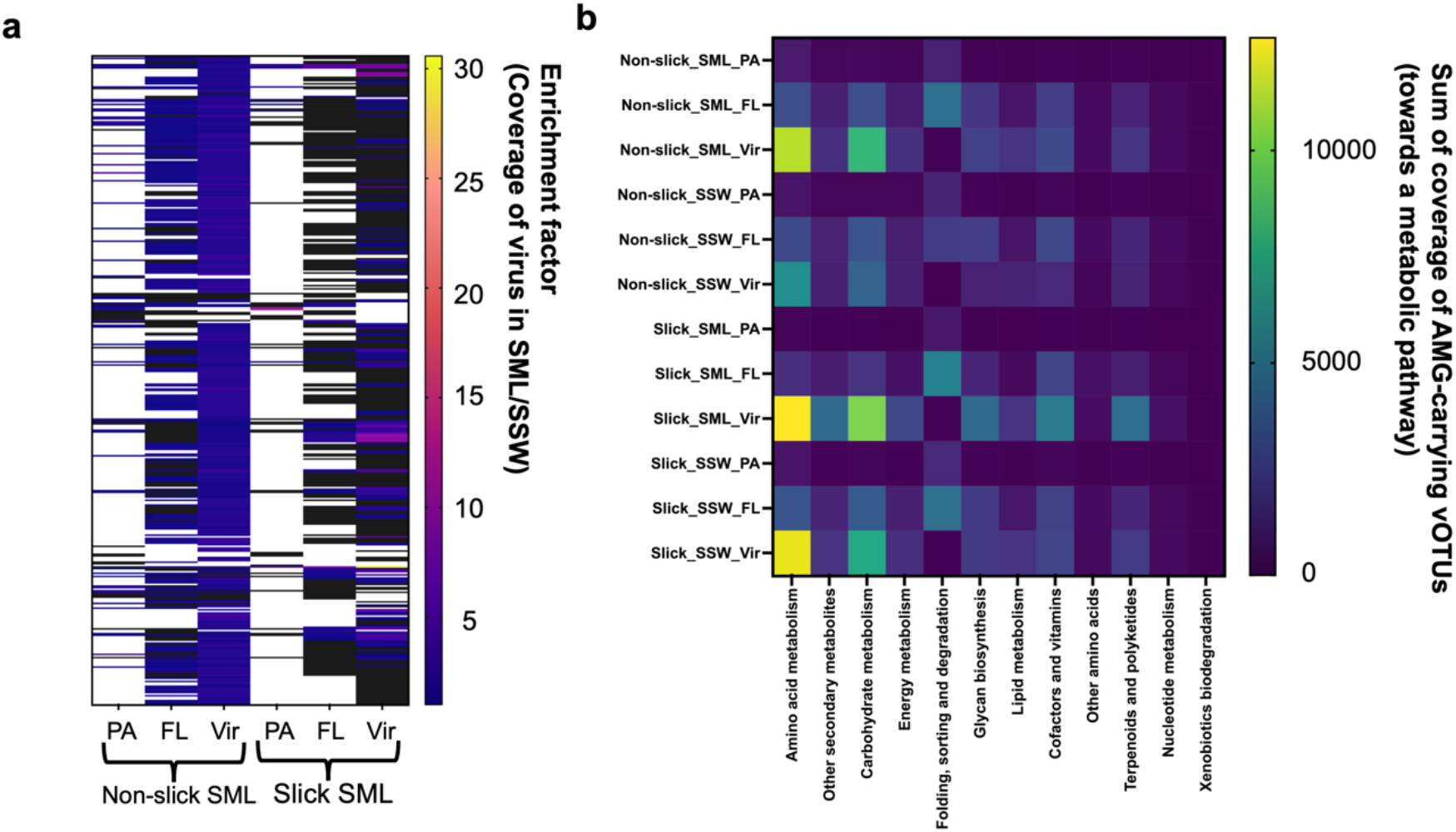
Viral enrichment in slick versus non-slick SML and viral auxiliary metabolic genes. Enrichment of viruses in slick SML and non-slick SML across different filtered fractions. Shown are coverage ratios > 1 indicating enrichment of a virus in the SML compared to the corresponding reference subsurface water sample. Black areas indicate depletion of a virus (ratio < 1), while white areas show absence of the virus in the nominator or denominator of the ratio. Values and VCs of the heatmap’s y-axis are given in Table S8a **(a)**. Coverage of vOTUs carrying an auxiliary metabolic gene (AMG) for KEGG categories **(b)**. Only vOTUs being present in a sample based on read mapping were considered for this analysis. Full information on involved AMGs is given in Table S8b. FL = free-living fraction (5 – 0.2 μm), PA = particle-associated fraction (> 5 μm), SML = sea-surface microlayer, SSW = subsurface water, Vir = viral fraction (< 0.2 μm)

### Virus-host predictions and slick SML specific CRISPR spacer-protospacer

Virus-host interactions were investigated based on shared protein clusters (VC information) with phages from known hosts, shared k-mer frequency patterns, and spacer-protospacer matches. The 34.1 kb and 40.7 kb vOTUs from slick SML were related to *Shewanella* phages based on protein-sharing network analysis (Table S6). These phages and an abundant 106.6 kb vOTU abundant in the FL fraction (463 x coverage) could be linked to diverse gammaproteobacterial MAGs based on shared k-mer frequencies, VC information, and spacer to protospacer matches (Fig. 7a, Table S6). Another 57.4 kb vOTU abundant in the virome (185 x coverage) was related to a *Pseudomonas* phage according to vConTACT2 (Table S6) but did not share k-mer frequency patterns with any of the MAGs. Two additional phages of 69.6 and 191.8 kb length detected only in slick SML had conflicting host evidence (Fig. 7a). MAGs from orders Flavobacteriales and Rickettsiales were k-mer linked to minimum 51 and 52 vOTUs, respectively, and likely represent hosts for viruses in the first 70 cm of the water column (Fig. S12).

**Fig. 7:**
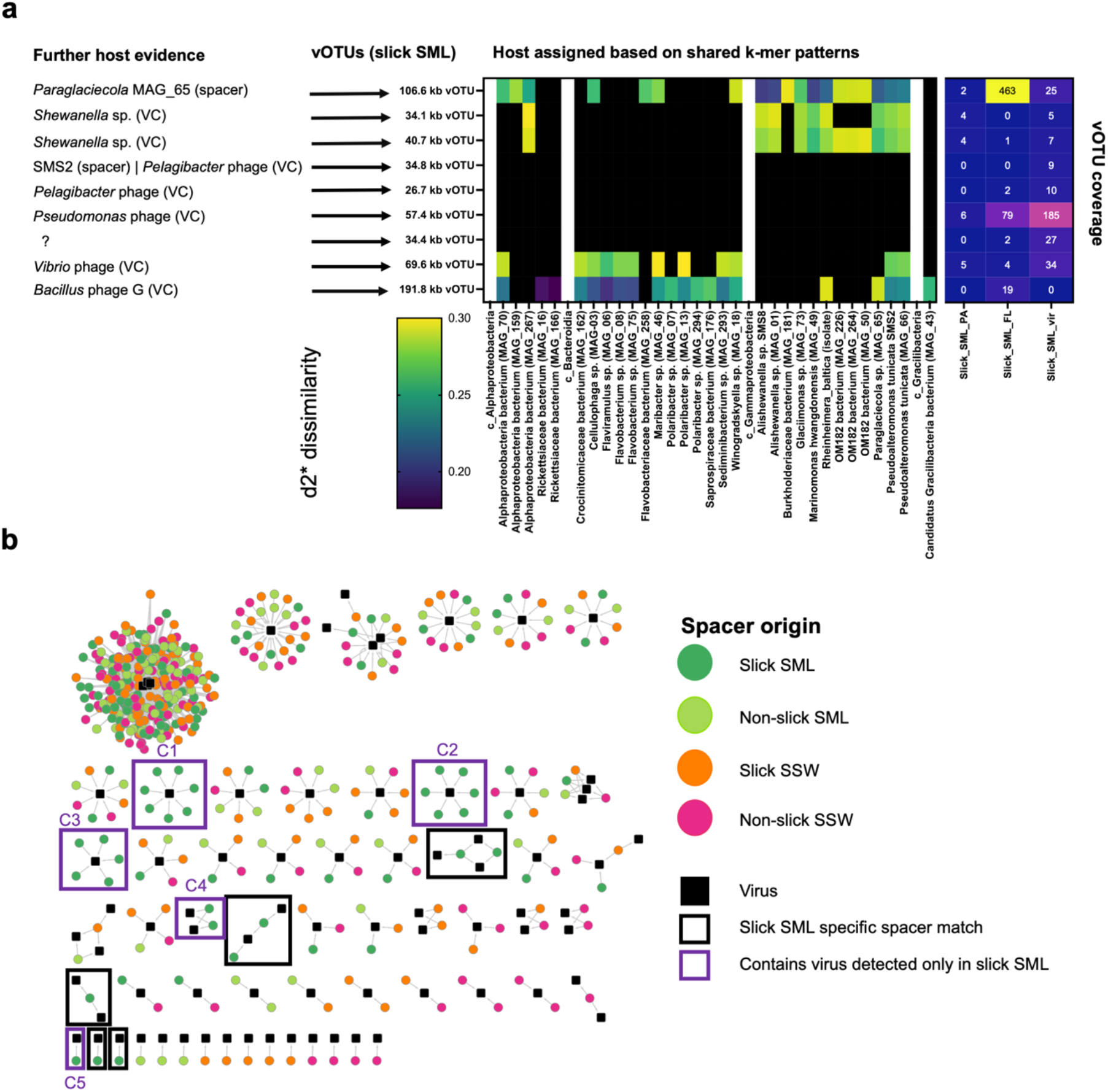
Slick SML-specific phages and their putative hosts. Based on k-mer frequency patterns, vOTUs were predicted to match host MAGs and isolate genomes (middle) and further host evidence (left) based on vConTACT2 viral clustering (VC) with known phages from reference database and CRISPR spacer matches from MAGs was consulted. Heat-map (right) depicts the coverage of vOTUs in the three size fractions. D2* is a dissimilarity measure (the lower, the higher the similarity) **(a)** CRISPR-spacer to vOTU protospacer matches at 100 % similarity reveal ten clusters with slick SML derived spacers, with C1-C5 including a slick SML specific vOTU from (a), framed in purple **(b)**.

Four MAGs comprised high-confidence CRISPR arrays (Table S4), but from these only a 32 bp spacer from the *Paraglaciecola* sp. (MAG_65) CRISPR array matched a protospacer of the highly abundant 106.6 kb vOTU exclusively detected in slick SML. In addition, we found another CRISPR spacer from a *Paraglaciecola* sp. MAG from slick SML matching 104 phages of the genus *Barbavirus* previously isolated on *R. baltica* [49,93] (Table S9). In addition, the genome of the bacterial isolate *P. tunicata* SMS2 and *R. baltica* SMS4 had a CRISPR array with a spacer of SMS2 matching a 34.8 kb vOTU only detected in the slick SML (Table S9; Supplementary Results).

Among CRISPR spacers recovered from metagenomic reads, 378, 326, 360 and 349 CRISPR spacer-protospacer matches at 100% similarity were detected in slick SML, non-slick SML, slick SSW, and non-slick SSW, respectively. These spacers originated from 29 different CRISPR arrays (Fig. S13). Slick SML-derived spacers targeted protospacers of six out of the nine slick SML specific vOTUs, namely the most abundant 57.4 kb (C1) and 106.6 kb (C3) vOTUs, and the 191.8 kb (C2), 34.1 kb, 40.7 kb (both C4), 69.6 kb (C5) vOTUs (Fig. 7, Table S10). Interaction clusters C1 – C4 contained CRISPR spacers originating from different CRISPR arrays/DR sequences (Fig. S13), suggesting infection histories with various bacteria. Blasting of the CRISPR DR sequences from those arrays revealed that bacteria hosting these arrays belonged to different Gammaproteobacteria, e.g., spacers from arrays P15 (C3), P16 (C3), P22 (C1), P40 (C1), P43 (C1) (Table S11, Fig. S13). In addition, one DR sequence had a Blast hit to Flavobacteria (P108, Table S12) and associated CRISPR spacers matching the virus were derived from all four sample types.

### Three phages lytic for *Alishewanella* sp. and *P. tunicata* extracted from slick SML

Two phages with myovirus morphology with lytic activity against *Alishewanella* SMS8 were isolated from slick SML at sites #1 and #3, and one phage with podovirus morphology for *P. tunicata* from slick SML at site #1 (Fig. 8a). Phages were named *Alishewanella* phage vB_AspM_Slickus01, vB_AspM_Slicko01, and *P. tunicata* phage vB_PtuP_Slicky01 in accordance with Kropinski, et al. [94] (further referred to as Slickus, Slicko, and Slicky). Slickus and Slicko had a mean head diameter of 85.8 ± 2.9 nm (n = 11) and 84.0 ± 2.6 nm (n = 15), and a mean tail length of 126.6 ± 3.5 nm (n = 8) and 126.1 ± 8.1 nm (n = 12), respectively. Slicky’s head diameter was 65.2 ± 2.5 nm (n = 5) (Table S12). Due to intergenomic similarity of 91.4 % in VIRIDIC for the *Alishewanella* sp. phages, we here propose the new genus Alishvirus with species names Alishvirus slickus (for strain Slickus) and Alishvirus slicko (for strain Slicko). For Slicky, we propose genus and species names Pseutunvirus and Pseutunvirus slicky, respectively. The genomes of Slickus, Slicko, and Slicky had a length of 141644, 141431, 66316 bp with 195, 198, and 81 open reading frames, respectively, and all carried transfer RNAs. Read mapping showed that Slickus and Slicko were only detected in the slick SML, while Slicky was below detection limit in all metagenomes (Fig. 8b). However, all phage isolates were targets of CRISPR spacers isolated from slick SML, and the SMS8 prophage additionally by spacers from non-slick SML but not from SSW (Fig. 8c). Slickus and Slicko had limited protein similarities with *Pseudomonas* phage pf16 and two *Rheinheimera* Barbaviruses while Slicky was related to *Vibrio* phage CHOED but not much to other known *Pseudoalteromonas* phages (Fig. 8d). Functional annotations of the genomes showed that Slickus and Slicko encode for structural proteins (tail fiber, tube and sheath protein, prohead core protein, major head subunit, head completion protein, portal vertex of the head, neck protein), DNA processing, repair and replication proteins (DNA polymerases, ligase, helicase, recombination endonucleases, DNA adenine methylase, putative polynucleotide 5’-kinase and 3’-phosphatase), proteins related to nucleotide metabolism (thymidylate synthase, ribonucleoside diphosphate reductase, ribonucleotide reductase, tRNA nucleotidyltransferase), biosynthetic enzyme (nicotinate phosphoribosyltransferase) and catalytic proteins (metallopeptidase). Slicky contained genes for structural proteins (coat protein), DNA processing, repair and replication proteins (DNA polymerase, DNA primase, endonuclease, 5 - 3 exonuclease), DNA packaging (terminase, portal protein), nucleotide metabolism (ribonucleotide reductase, ribonucleoside-diphosphate reductase, thymidylate synthase), biosynthetic enzymes (nicotinate phosphoribosyltransferase, dimodular nonribosomal peptide synthetase), repair enzymes (glutamine transaminase) and catalytic proteins (lysozyme, gamma-glutamylaminecyclotransferase) (Table S13). To our best knowledge, these are the first phage isolates reported for these host species. Slickus, Slicko, and Slicky tested negative for cross infections in a host range experiment involving other slick SML bacterial isolates (see Supplementary Results).

**Fig. 8:**
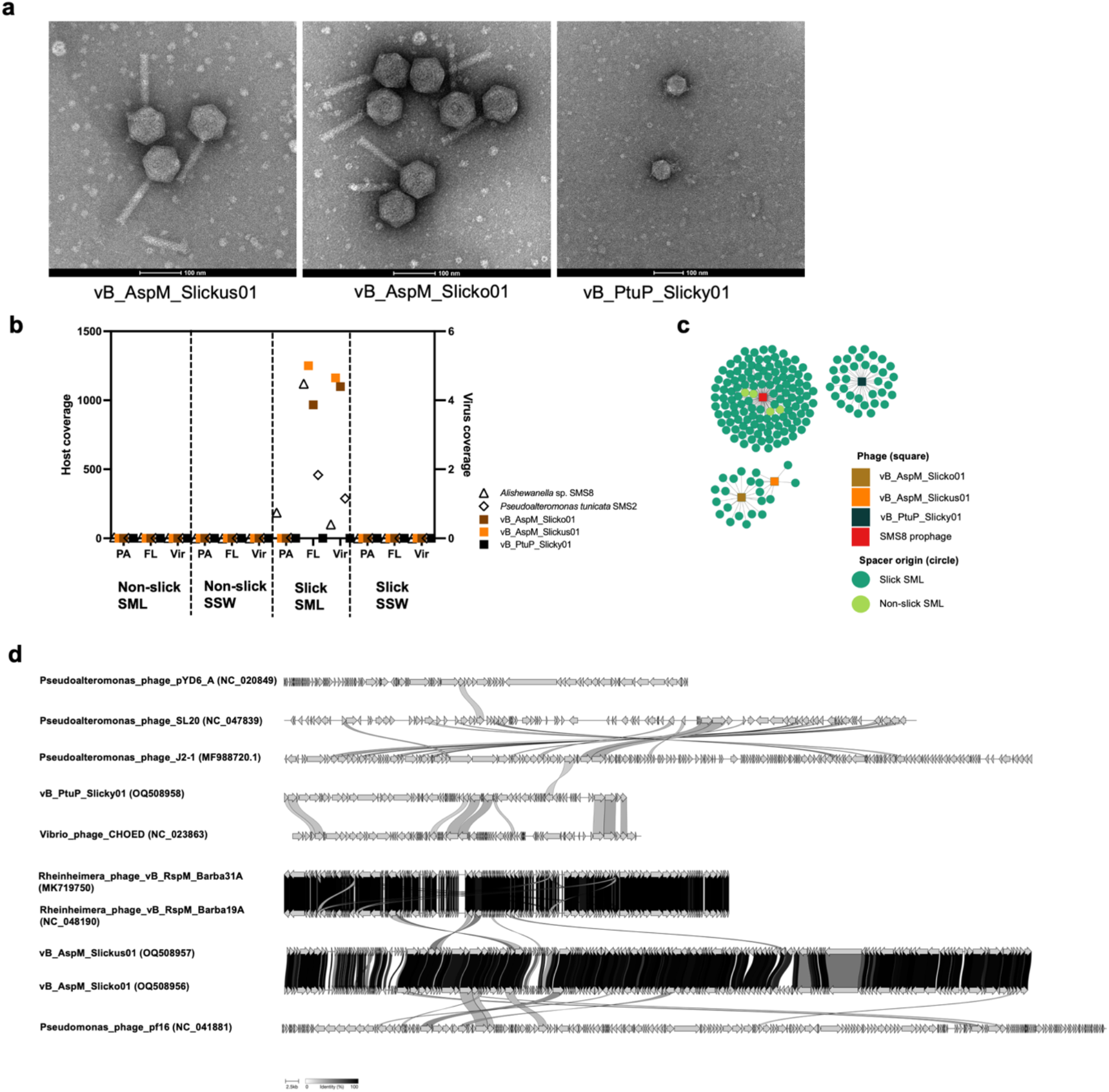
Transmission electron microscopy images, abundance, CRISPR spacer matches and synteny of lytic phage isolates extracted from slick SML. Negatively stained electron micrographs reveal myovirus morphology of Slickus and Slicko and podovirus morphology of Slicky, scale bar 100 nm **(a)**. Coverage of reads based on mapping to the host MAGs Alishewanella sp. (MAG_01) and Pseudoalteromonas tunicata (MAG_66) as well as phages vB_AspM_Slickus01, vB_AspM_Slicko01, and vB_PtuP_Slicky01 **(b)**. CRISPR spacer extracted from reads matching at 100 % similarity to phage genomes from isolates and the SMS 8 prophage of 50 kb. No matches from SSW spacers were detected. **(c)**. Synteny plots of Slickus, Slicko, and Slicky genomes with indicated identity of proteins. Annotations for the phage isolates are given in Table S13 **(d)**. SML = sea-surface microlayer, SSW = subsurface water

## Discussion

In this work, we provide new insights on viral-bacterial diversity and interactions in the SML of a natural surface slick, which forms under calm conditions on water surfaces. Metagenomic analysis revealed distinct viral and bacterial communities with specific functional properties in the SML of this wave-dampened zone in comparison to non-slick SML and SSW.

Our study distinguishes two types of bacteria and viruses: 1.) “surface cosmopolitans”, *i.e*. bacteria and viruses that are present in all sample types and might be transferred into the slick with rising bubbles [95], and 2.) “slick-specific opportunists”, *i.e*. bacteria and viruses being detected in slick SML only and responding quickly to emerging slick conditions.

Group 1 comprised a VC from slick SML related to flavobacterial phages as well as the many vOTUs matching to *Flavobacteriaceae* and *Rickettsiaceae* MAGs based on k-mer analysis. Flavobacterial phages often occur in timely proximity to phytoplankton blooms and heterotrophic bacterial communities following up on such blooms [96], for instance in three consecutive summers at the Linnaeus Microbial Observatory in the Baltic Sea [93]. In addition, we found several flavobacterial (*Flaviramulus* sp. (MAG_06), *Flavobacterium* sp. (MAG_08)) and an alphaproteobacterial MAG (*Rickettsiaceae* MAG_166) occurring in SML and SSW and representing hosts for these phages. A CRISPR array with DR sequence linked to Flavobacteria had spacers from SML and SSW samples matching different vOTUs indicating presence of this CRISPR system in both water layers. Common in the Baltic Sea, *Cellulophaga baltica* are hosts for diverse flavobacterial phages with varying degree of infection susceptibility [97,98], and *Cellulophaga* and *Polaribacter* were predicted as actively replicating in the FL fraction of the slick.

Group 2, typical for the slick SML, includes viruses infecting Gammaproteobacteria, some of which were detected exclusively in the slick SML, which is presumably because they were beyond detection limit in the other samples. The prevalence of Gammaproteobacteria in short-lived but particle-rich slicks [26] corresponds to a similar occurrence in ephemeral sea foams from the air-water boundary [11], and comparable virus-host ratios (>40) in foams and slicks [32]. During slick formation, which can happen within an hour [99], Gammaproteobacteria like *R. baltica, Alishewanella* sp., and *P. tunicata* can possibly respond quickly to labile carbon compounds as reported for other Gammaproteobacteria [100,101]. As motility-mediating chemotaxis genes in isolate genomes and mostly in MAGs with high abundance in slick SML were found, we speculate that these bacteria actively migrate towards and/or within the slick to reach nutrients. Presence of GH103 and GH171 in *R. baltica* further suggest the ability to degrade peptidoglycan from detrital bacterial matter, so that this strain could particularly benefit from high turnover of cells in the slick. In addition, *R. baltica* and *Alishewanella* sp. contained homoserine lactone genes, whose presence is typical for biofilms [102] indicating that intraspecific communication is probably important in the slick SML. Contrasting CAZyme patterns of slick SML low and high abundant MAGs could show that different trophic niches are occupied by these groups. A similar pattern was observed for the *P. tunicata* and *R. baltica* isolates, potentially explaining their successful co-existence in the slick. High abundance of *Alishewanella* sp. in the PA fraction indicates a surface-associated lifestyle and biofilm-forming abilities, supported by the observation of biofilms in flask cultures of SMS8 and SMS9. *P. tunicata* favors a surface-associated lifestyle in the marine environment [103], making both bacteria ideal candidates for inhabiting the SML featuring biofilm-like properties [26].

Gammaproteobacterial hosts from slick SML possess a slick-specific repertoire of CRISPR spacers to interact with slick-specific vOTUs, and also *P. tunicata* SMS2 contained a CRISPR array, a known feature for this species [103]. Remarkably, the same slick vOTU was targeted by spacers from different CRISPR arrays, which due to the conservation of DR sequences [104] likely belong to different host strains. Potentially, abundant slick vOTUs have tried to infect multiple hosts within the class Gammaproteobacteria, despite such broad host ranges being rare in nature [105,106]. Protospacers could have been acquired from defective phages [107] or from intact phages without phage replication being necessarily successful. Possible is that a high viral micro-diversity enables different host ranges, or that the same protospacers exist in different viruses. Extensive spacer exchange might be enhanced by a higher likelihood for virus-host encounters through enhanced virus-host coupling in the neuston compared to the plankton [32].

Lytic infections between two abundant bacteria in the slick and three phage isolates were found, suggesting that virus-host dynamics in slick SML follow the kill-the-winner theorem [34,35]. The presence of a prophage in *Alishewanella* (Fig. S5) with no similarities to the lytic phages allows speculations that harsh environmental conditions at the air-sea interface occasionally favor the prophage state, or that lysogeny is favored when the bacterium leaves the air-sea interface upon slick disintegration. Interestingly, slick SML VCs of the PA fractions carried genes for reverse transcriptase, which have been previously found to predominate in marine plankton assemblages in the > 5 μm size fraction as part of eukaryotic retrotransposons or retroviruses [108]. Retroviruses can infect oceanic eukaryotes (reviewed by Lang, et al. [109]). This shows that eukaryotic viruses were part of the studied viral community, but due to our study focus on phages, their role in slick SML and interactions with hosts remains to be elucidated.

Conditions at the air-sea interface and the different physicochemical environment (high surfactant loads, low surface tension) compared to SSW likely contributed to the distinct viral-bacterial community structures in the slick. Bacteria like *Pseudomonas* or *Pseudoalteromonas* are known surfactant and exopolysaccharide producers [110], respectively. Slick SML dwelling *P. tunicata* encoded gene homologs for biosurfactant production. Surfactants could contribute to lowering surface tension and hence facilitate surface colonization, but surfactin also exerts anti-viral properties [111], suggesting that surfactant-phage interactions in slicks require further research. Light conditions could be an additional factor for shaping viral-bacterial communities in slicks due to the varying responses of *Oceanospirillales, Pseudoalteromonas* and Flavobacteria to light, dependent on the particle attachment status [112], but was not considered in this study. The SML is strongly affected by diel light effects [113], and bacteria and likely their phages have to adapt to solar and ultraviolet radiation with one light adaptation posing the typical pigmentation of SML bacteria [3,114]. This matches phenotypic observations of purple-blue coloring in *Alishewanella* SMS9, all *R. baltica* strains and *P. tunicata* SMS2 with pigment violacein being responsible for coloration of the latter (Table S2f) as previously reported for this species [103].

## Conclusions

Our results support that the slick SML harbors distinctive viral-bacterial communities with adaptations to a habitat that has a short lifespan and forms quickly thus posing exceptional colonization demands on its inhabitants. Gammaproteobacteria in slick SML showed functional adaptations such as pigmentation, prevalence of genes related to chemotaxis and quorum sensing, higher diversity of CAZymes, biofilm formation as well as adaptive immunity targeting specific slick phages. Despite experiencing a diversity drop in the slick, phages infected typical and abundant slick bacteria and different vOTUs became enriched, with proliferation being likely a strategy to increase the chance for point mutations to circumvent attacks by the host’s adaptive immunity. We conclude that virus-host interactions align to the peculiarity of slicks and uncouple from remaining waters. High selective pressure exerted by prevailing conditions and an active virus-host arms race might favor discovery of previously unknown viruses and bacteria as shown in this work. Viral lytic activity combined with a high DOC level suggest that slicks can be hotspots for the viral shunt at the air-sea interface, representing an overlooked factor for carbon cycling in the sea.

## Supporting information

Suppplementary Tables

## Acknowledgements

JR received funding for the project “Exploring the virioneuston: Viral-bacterial interactions between ocean and atmosphere (VIBOCAT)” by the German Research Foundation (DFG RA3432/1-1). We acknowledge funding for sequencing by the Anna-Greta and Holger Crafoord Foundation for the project “The impact of viruses on bacterial biodiversity investigated through time-series metagenomics” (CR2019-0034). HAG was funded by the DFG, grant/award no. 34509606: Ökologie, Physiologie und Molekularbiologie der Roseobacter-Gruppe: Aufbruch zu einem systembiologischen Verständnis einer global wichtigen Gruppe mariner Bakterien.

We like to thank Carola Lehners for technical assistance as well as Thomas Mollica and Emil Fridolfsson for advice regarding DOC sampling. We further thank Samuel Hylander for lending a Ruttner sampler, and Hannah Schweitzer for advice with biosurfactant genes. Data handling was enabled by resources provided by the Swedish National Infrastructure for Computing (SNIC2020-16-49) at UPPMAX partially funded by the Swedish Research Council through grant agreement no. 2018-05973. We like to thank Pavlin Mitev at UPPMAX for support.

## Competing Interests

The authors have no competing interests to declare.

## Data Availability Statement

Cell and VLP abundance, surfactant and DOC data were submitted to PANGAEA data publisher under https://doi.pangaea.de/10.1594/PANGAEA.955904 [115]. Sequence data for this project are stored in Bioproject PRJNA855638, physical Biosamples SAMN29538895 – SAMN29538900 and SAMN29499491 – SAMN29499496 at NCBI. Raw reads of samples are stored in sequence read archive (SRA) under accession numbers SRR20006720 – SRR20006725 and SRR20314586 – SRR20314591. Metagenome-assembled genomes (316) correspond to Biosamples SAMN29881236 – SAMN29881551 and accession numbers JANQTE000000000 – JANQZZ000000000 and JANRAA000000000 – JANRFH000000000. The viral metagenome of 428 scaffolds is stored within Biosample SAMN31653011, accession number JAPJDW000000000. Genomes of bacterial isolates are stored within Biosample SAMN31710611 – SAMN31710617, accession numbers JAPJDX000000000 – JAPJED000000000. Genomes of phage isolates Slickus, Slicko, and Slicky are stored at GenBank under accessions OQ508956 – OQ508958. Data will be released upon peer-reviewed publication of the manuscript. For further information, please see Table S14.

## Notes

### Competing Interest Statement

The authors have declared no competing interest.

