## Supplementary Material for "Ecogenomics reveals distinctive viral-bacterial communities in the surface microlayer of a natural surface slick"

**List of Supplementary tables (provided as Excel file):**

Table S1: Sampling notes

Table S2a: Bacterial strains from slick SML isolated and sequenced in this study

Table S2b: Similarities of bacterial slick SML isolates to type strains based on digital DNA-DNA hybridization determined using the Type Strain Genome Server (https://tygs.dsmz.de)

Table S2c: KEGG numbers assigned to genes from slick SML isolates

Table S2d: Metabolic pathways in bacterial slick SML isolates, as reconstructed from predicted KEGG numbers.

Table S2e: Predicted genes encoding carbohydrate-active enzymes (CAZymes) in bacterial slick SML isolates.

Table S2f: Predicted biosynthetic genes in bacterial slick SML isolates.

Table S2g: Predicted biosurfactant-encoding genes in bacterial slick SML isolates.

Table S3a: CRISPRcasFinder (Couvin et al., 2018) output about evidence level 4 CRISPR arrays and associated consensus DR sequences

Table S3b: List of consensus direct repeat (DR) sequences used for spacer extraction and CRISPR array number they refer to

Table S4a: Information on metagenome-assembled genomes (MAGs), inlucinding assigned taxonomy, completeness, contamination, CRISPR repeat, CRISPR spacer to protospacer matches

Table S4b: MAG grouping based on high abundance in slick SML (Slick_highAb), low abundance in slick SML (Slick_lowAb), and high abundance in the underlying seawater (SSW).

Table S4c: KEGG numbers with significantly different fractions between MAG groups, displayed as relative fraction of all genes. Coloring illustrates the group with highest, significantly enriched fraction.

Table S4d: Predicted genes encoding carbohydrate-active enzymes (CAZymes) in MAGs.

Table S4e: Predicted biosurfactant-encoding genes in MAGs.

Table S5: Coverage output by iRep for dereplicated MAG set and SMS3

Table S6: CheckV (Nayfach et al., 2020) information on viral OTUs andviral clusters (VC) assigned by vConTACT2 (Bolduc et al., 2017) and compiled by graphanalyzer

Table S7: Virus enrichment (enrichment factor>1) data corresponding to Fig. 6A. EF is a ratio of coverage in an SML sample divided by SSW counterpart. If a value is missing, it means that either the virus was absent in SML, SSW or both or that the value was 0.

Table S8a: Sum of coverage of viruses carrying an auxiliary metabolic gene (AMG) towards a certain pathway.
Table S8b: Info of AMG (KEGG orthology) detected on viral OTU; Only class I AMGs (Hurwitz et al., 2015) were considered.
Table S9: CRISPR spacer (from MAGs and slick SML bacterial isolates) to protospacer matches of Baltic Sea phage isolates and the viral populations from this study.
Table S10: Spacer matches to vOTUs at 100% similarity.
Table S11: Blastn results for direct repeat (DR)/array sequences shown in Fig S13.
Table S12: Head and tail measurements of phage isolates according to protocol Brum (2011).
Table S13: Functional annotations/output of DRAMv (Shaffer et al., 2020) for 428 vOTUs, lytic phage isolates and Alishewanella sp. (SMS8) prophage
Table S14: Accession number for Bioproject PRJNA855638: Prokaryotes and viruses from sea-surface microlayer of a brackish surface slick

**Supplementary results:**

**Host range experiment**

Plaque assays of the two lytic *Alishewanella* phage vB_AspM_Slickus01 and vB_AspM_Slicko01 and *Pseudoalteromonas tunicata* phage vB_PtuP_Slicky01 were performed on host strains SMS2, SMS3, SMS4, SMS8, SMS9, SMS11, and SMS12. This was conducted to test for cross-infection due to the phylogenetic relatedness of *Alishewanella* sp., *P. tunicata* and *Rheinheimera baltica* and due to the observation of spacers from different Gammaproteobacteria targeting the same vOTU, and spacers from gammaproteobacterial MAGs such as *Paraglaciecola* having spacer to protospacer matches to Barbaviruses that were isolated on *R. baltica.* In addition, Slickus, Slicko and Slicky possess tRNAs, and tRNAs can have a role in cross-infectivity, at least for cyanobacterial hosts [1].
Overnight cultures of the bacterial strains were used for the plaque assays. Bacteria (300 µl) were mixed with 3.5 ml top-agar and spread on Zobell agar. Ten-times dilutions of phages until $10^{-7}$ were produced in 1.5 ml tubes and 10 µl were spotted on Zobell agar plates containing the solidified bacteria and top-agar mix. Clearing of the bacterial lawn by phage lysis (plaques) or

bacterial inhibition was investigated after 48 hours. As a result, we did not find any cross-infection in a host range assay where the phages were tested on the above-mentioned bacterial isolates. *Alishewanella* phage vB_AspM_Slickus01 and vB_AspM_Slicko01 only infected *Alishewanella* sp. SMS8, and vB_PtuP_Slicky01 only *P. tunicata* SMS2.

**Relative abundance (Fig. 2)**

The family *Chromatiaceae* comprises *Alishewanella* sp. and *R. baltica*, which are phylogenetically closely related species [2]. Since *Alishewanella* sp. from the slick SML is likely a new species (see below) with no reference genome in the database of the tool we used for taxonomic profiling, *Alishewanella* sp. has been overseen or falsely taken as *R. baltica*, as from 17.8 % relative abundance of *Chromatiaceae*, 17.1 % were *R. baltica*, although *Alishewanella* sp. was ~10 x more abundant based on further investigations (see main text, Table S5). This is the reason why we report taxonomy at family level in this case.

**CRISPR system of *P. tunicata* SMS2:**

Two different evidence level 4 CRISPR arrays associated with two different consensus DR sequences (Table S2a) were detected on *P. tunicata* SMS2 by using CRISPRcasFinder [3]. The DR sequence of Array 1 "GTGAACTGCCGAGTAGGCAGCTGAAAAT" equipped with 10 spacers and Array 2 "TTTCTAAGCTGCCTGTGCGGCAGTGAAC" equipped with 36 spacers corresponded to DR186/P64 and DR54/P26 recovered from assemblies (Table S3b), respectively. Array 2 (position 78-2265 nt) was flanked by four genes, namely cmr6, cmr5, cmr4 and cmr1 of the Cas RAMP module (Cmr) effector complex belonging to Cas Type IIIB system and were detected on the same scaffold (position 3770-10512 nt) as Array 2. On three other scaffolds, Cas clusters for Cas3 (4 genes) and Cas3a (2 genes) of a Type I CRISPR system were found. Thomas, et al. [4] reported only one consensus DR sequence "GTTCACTGCCGCACAGGCAGCTCAGAAA" for *P. tunicata* in conjunction with Cas 1, 3 and 4 (typical of Type I CRISPR systems) as well as two unknown Cas homologs. The DR sequence of [4] differs only by one point mutation from the Array 2 DR sequence in reverse complement. Spacers extracted from metagenome reads based on the DR sequence from Array 2 "TTTCTAAGCTGCCTGTGCGGCAGTGAAC" (P26) and matching them to the virome of 428 OTUs revealed spacer matches to the two *Shewanella sp.* phage 1/41-related 34.1 kb vOTU and 40.7 kb vOTU in the slick SML, while a spacer recovered from Array 1 of *P. tunicata* SMS2 matched a 34.8 kb vOTU from the slick SML, which was related to *Pelagibacter* phage HTVC023P based on shared protein clusters (Table S6). Our results support that CRISPR systems for slick SML derived *P. tunicata* SMS2 are more complex than previously reported by using the Cas machinery of different CRISPR system types, and because the CRISPR-Cas Type III B system can use both DNAs and RNAs as substrates for spacer acquisition (reviewed by Zhang and An [5]). Type III systems are rather uncommon in Gammaproteobacteria (reviewed by Makarova, et al. [6]).

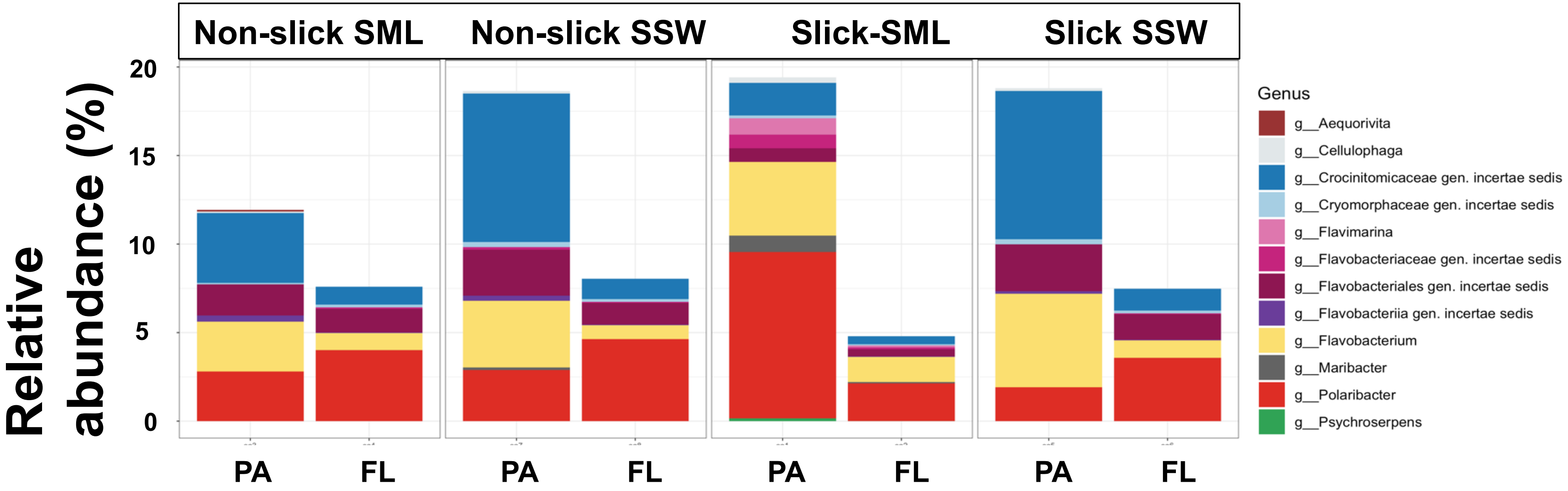


***Fig. S1:*** *Relative abundance of class Flavobacteriia based on results from mOTUs. FL = free-living fraction (5 - 0.2 µm pore size filtered), PA = particle-associated fraction (> 5µm filtered), SML = sea-surface microlayer, SSW = subsurface water.*

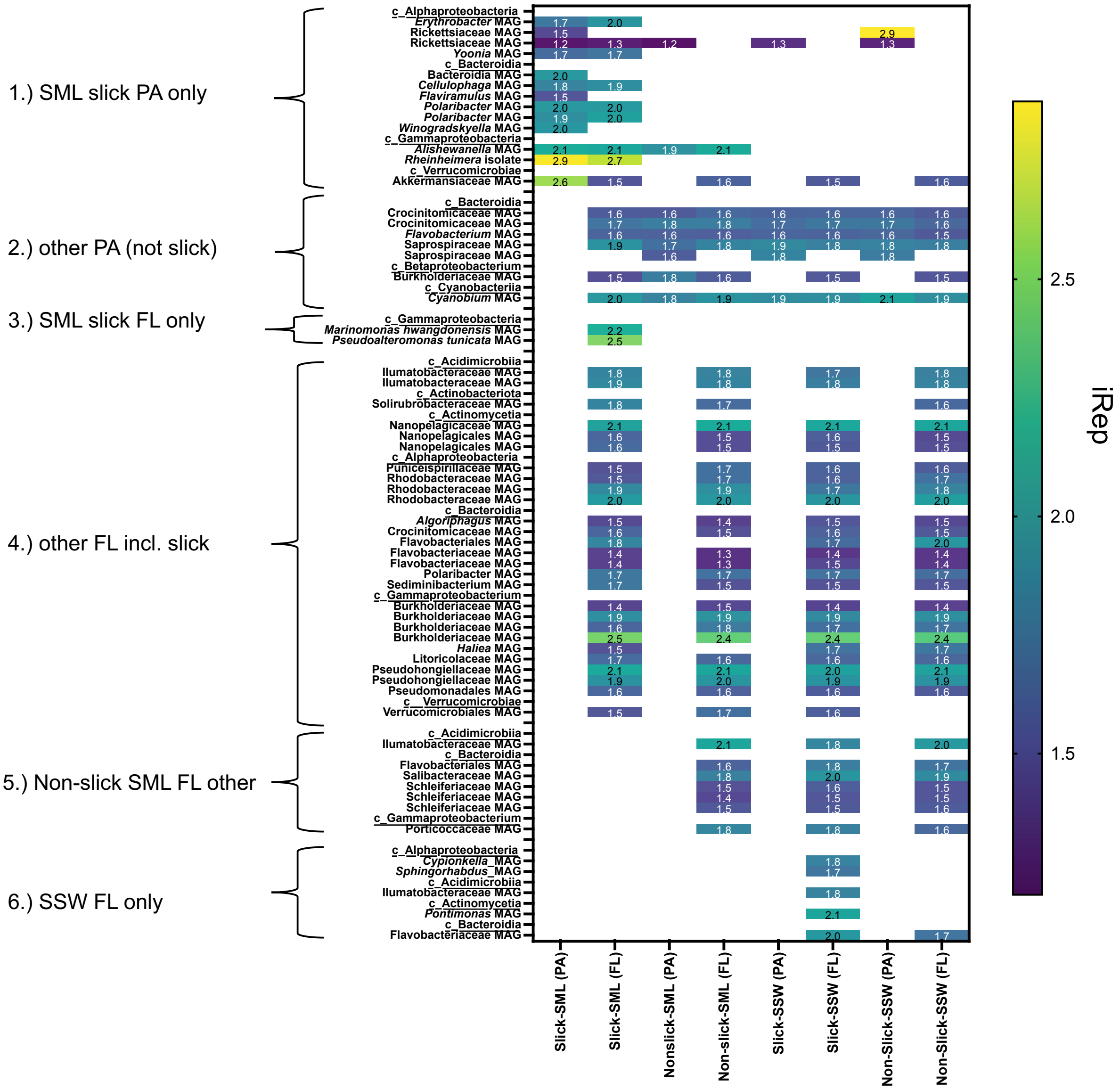


***Fig. S2:*** *Groups of actively replicating bacteria based on index of replication (iRep). We defined six groups according to samples, in which replicating bacteria were detected, e.g. "SSW FL only" contains MAGs with iRep in the SSW FL fraction only. Blank fields indicate that iRep values could not be predicted within the defined thresholds. An iRep value of 2 indicates that the coverage at the origin of replication is double the coverage at the terminus. This could be achieved if half of the population was in process of two simultaneous replication events or each cell of the population had started one replication. FL = free-living fraction (5 - 0.2 µm pore size filtered), PA = particle-associated fraction (>5µm filtered), SML = sea-surface microlayer, SSW = subsurface water*

**a**

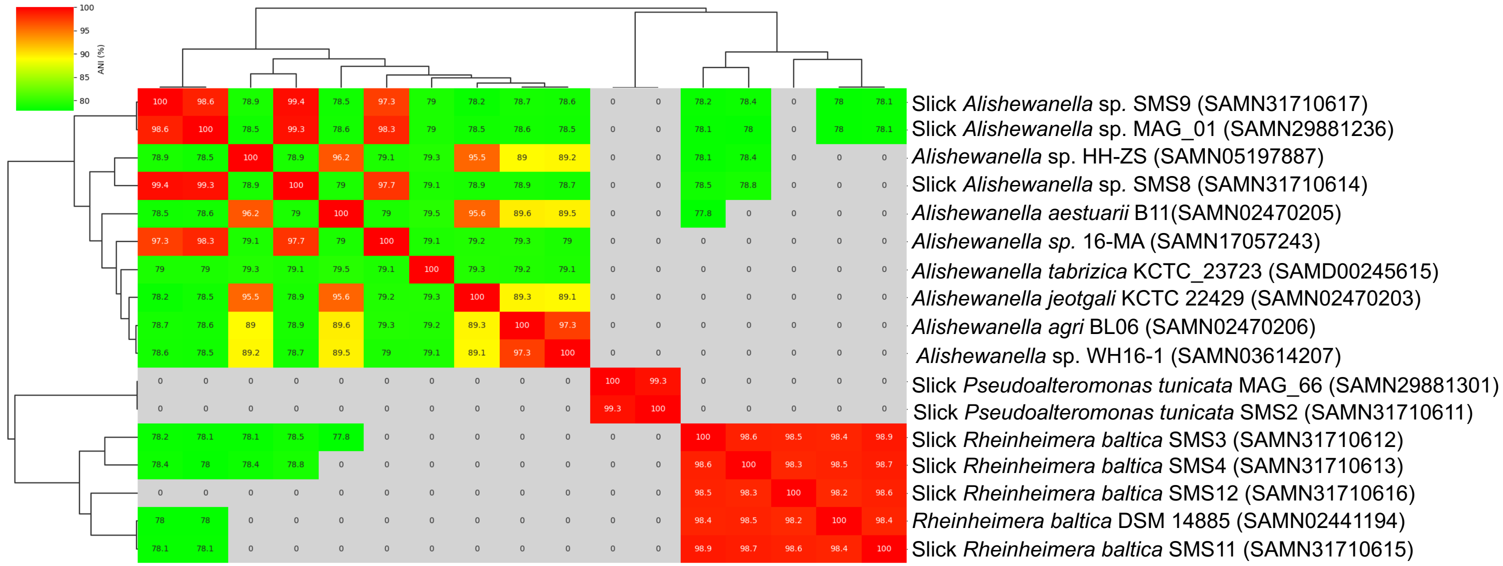


**b**

| **Average nucleotide identity (ANI) and aligned fraction to Slick *Alishewanella* sp. SMS8 (SAMN31710614)** | | | | |
|---|---|---|---|---|
| **Genome** | **ANIb [%]** | **Aligned [%]** | **Aligned [bp]** | **Total [bp]** |
| Slick *Alishewanella* sp. SMS9 (SAMN31710617) | 99.39 | 58.43 | 2086875 | 3571747 |
| Slick *Alishewanella* sp. MAG_01 (SAMN29881236) | 99.16 | 84.74 | 3026788 | 3571747 |
| *Alishewanella* sp. 16-MA (SAMN17057243) | 97.24 | 88.91 | 3175681 | 3571747 |
| *Alishewanella tabrizica* KCTC_23723 (SAMD00245615) | 74.08 | 61.9 | 2211085 | 3571747 |
| *Alishewanella* sp. WH16-1 (SAMN03614207) | 72.92 | 59.61 | 2129233 | 3571747 |
| *Alishewanella agri* BL06 (SAMN02470206) | 72.88 | 60.48 | 2160152 | 3571747 |
| *Alishewanella* sp. HH-ZS (SAMN05197887) | 72.74 | 58.26 | 2080864 | 3571747 |
| *Alishewanella aestuarii* B11(SAMN02470205) | 72.7 | 58.29 | 2082122 | 3571747 |
| *Alishewanella jeotgali* KCTC 22429 (SAMN02470203) | 72.69 | 58.91 | 2104109 | 3571747 |
| Slick *Rheinheimera baltica* SMS12 (SAMN31710616) | 71.04 | 47.5 | 1696542 | 3571747 |
| Slick *Rheinheimera baltica* SMS4 (SAMN31710613) | 71.01 | 52.13 | 1862029 | 3571747 |
| Slick *Rheinheimera baltica* SMS11 (SAMN31710615) | 70.85 | 51.87 | 1852563 | 3571747 |
| Slick *Rheinheimera baltica* SMS3 (SAMN31710612) | 70.84 | 51.78 | 1849277 | 3571747 |
| *Rheinheimera baltica* DSM 14885 (SAMN02441194) | 70.75 | 50.21 | 1793325 | 3571747 |
| Slick *Pseudoalteromonas tunicata* SMS2 (SAMN31710611) | 66.47 | 26.92 | 961342 | 3571747 |
| Slick *Pseudoalteromonas tunicata* MAG_66 (SAMN29881301) | 66.34 | 26.21 | 936297 | 3571747 |

***Fig. S3:*** *Slick SML Alishewanella sp. form a distinct bacterial cluster.* *Average nucleotide identity (ANI) comparison for Rheinheimera baltica isolates, Pseudoalteromonas tunicata isolate, Alishewanella sp. isolates and MAG assigned to Alishewanella including relevant reference genomes with Biosample number mentioned alongside. As FastANI does not output ANI much smaller than 77%, we investigated further for the comparison of Alishewanella sp. SMS8 and R. baltica and P. tunicata strains using ANIb analysis from JWSpeciesWS web server [7] with results and alignment fractions reported in the table.*

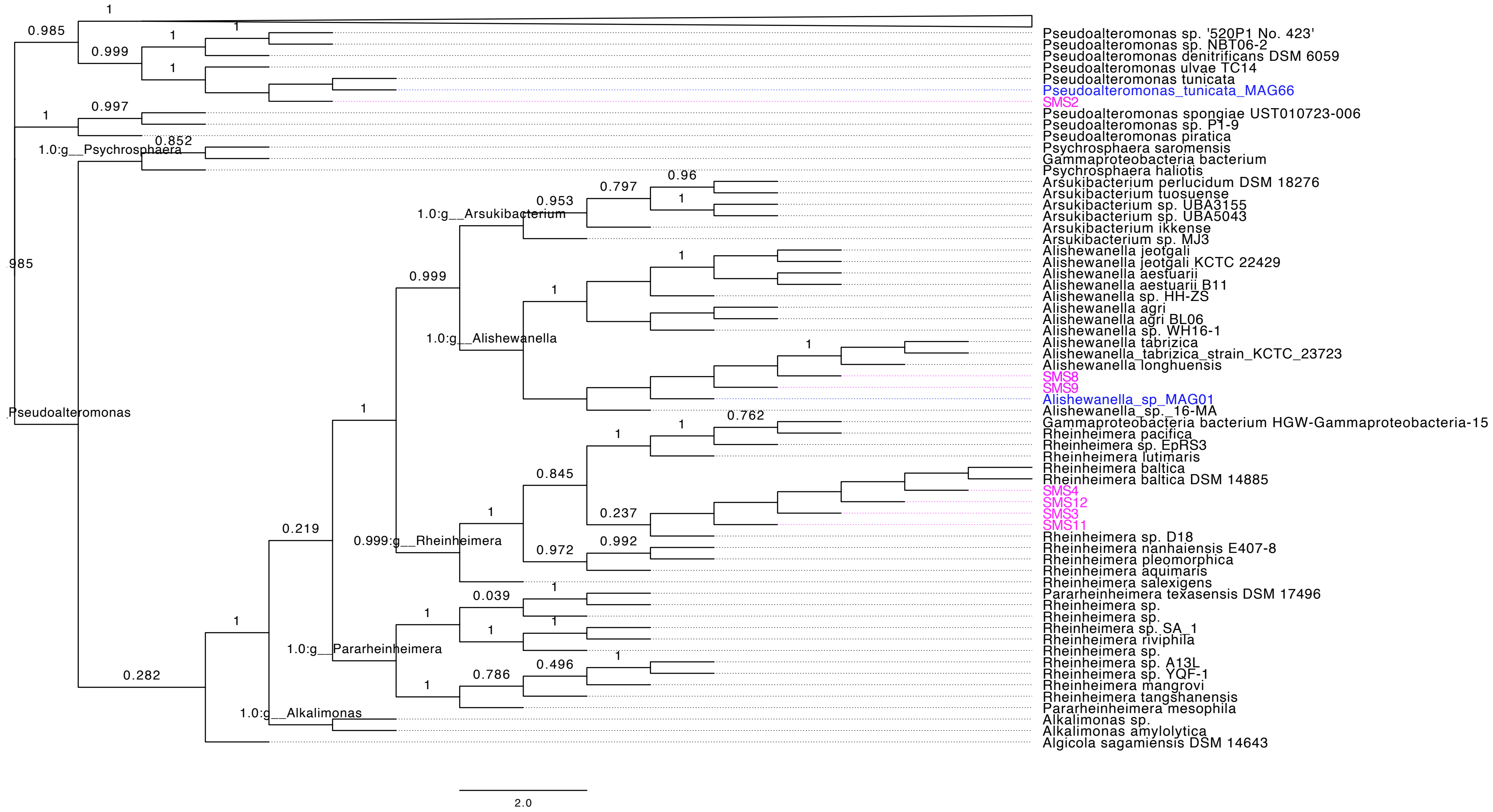


***Fig. S4:*** *Phylogenetic relatedness of SML isolates (pink) and respective MAGs (blue) within the bac120.classify.tree (identification uses 120 bacterial marker genes) predicted by the classify_wf in GTDB-Tk [8] (settings explained in the main text), which uses pplacer v.1.1 [9] to find the maximum-likelihood placement of genomes in the tree. A subnetwork of the full tree was extracted in Dendroscope v.3.8.5. [10], and the tree was rooted at the midpoint and tips aligned in FigTree v.1.4.4. [11].*

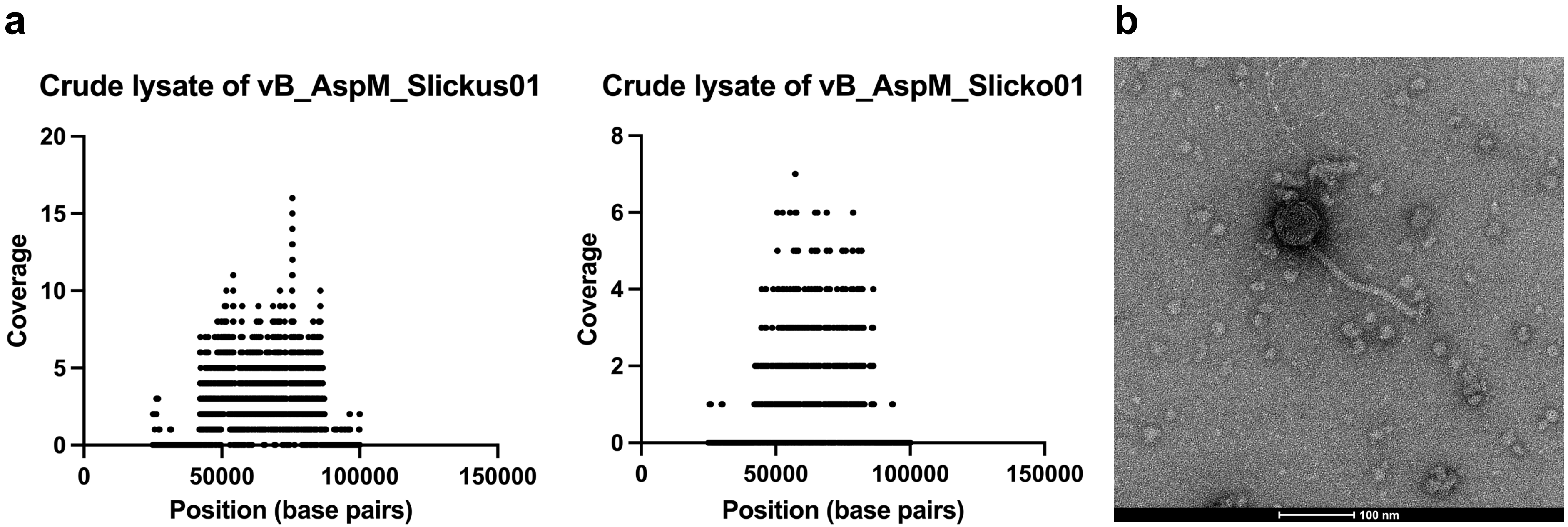


***Fig. S5:*** *A 50 kb-prophage from the Alishewanella sp. isolate SMS8 positioned between 36371 and 86348 bp in the genome's scaffold shows induction (high coverage) based on mapping of reads from the crude lysates of Alishewanella phage vB_AspM_Slickus01 and vB_AspM_Slicko01 to the prophage-containing scaffold of SMS8* ***(a)****. In transmission electron microscopy, a siphovirus structure among the lytic myovirus-like morphologies in the crude lysate of vB_AspM_Slicko01 was detected, potentially representing the 50 kb-prophage* ***(b)****.*

**Additional results prophage:**

The 50 kb prophage carries a tape tail measure protein of 1178 bp length (see below). According to Hoetzinger, et al. [12], the amino acid sequence length of the tape measure protein can be converted to tail length using the formula y= 14 + 0.144 * x, where x is the length of the tape measure protein (number of amino acids), which in case of the 50 kb prophage converts to a tail length of 183.6 nm. ImageJ gives a tail length of 169.4 nm for this prophage, which is reasonably comparable to the calculation. According to the linear regression presented by Hoetzinger, et al. [12], the chance of a phage having siphovirus morphology with a tape measure sequence > 1000 bp is high. Most prophages have siphovirus morphology, and the capsid diameter of ~65 nm is also consistent with the genome size of 50 kb according to [13]. The prophage has 68 open reading frames. Functional annotations of the genes revealed presence of lysogeny related proteins (two integrases, regulatory CII family protein), structural proteins (baseplate, two capsid proteins, tape measure protein), DNA processing, repair and replication proteins (single-strand DNA-binding protein, exonuclease, DNA replication protein DnaC, transcriptional repressors), DNA packaging (terminases, portal protein), nucleotide metabolism (thymidylate synthase) and catalytic proteins (lysozyme, permuted papain-like amidase) (Table S13). The prophage got targeted by CRISPR spacers extracted from reads of the slick SML and the non-slick SML sample, but not from the SSW (Fig. 8C, main manuscript).

>NG_32195_SMS8_spades_3_length_559453_cov_168_fragment_1_53 rank: D; Tape measure protein [PF20155.2] (db=pfam)
MSVKQKFIDLVLRGKDLFSPTASAASDELKKLQAESKTTSEEMRKLEQAQAQVAKA
QGLELFAKQAELALAGAREEVTRLAREMDASDRPTKEQSEALKLATRSASQLQTEY
NKLQSQLSRSKTELQQSGVNTANLASEQDRLQREVKESANALNEKRTKLRELRSDM
DTTEKSTGKFGEGLRGLTTRLAAFAAAYVGINQLRSALTAIFTTGDKFEKLDIQLTGI
MGSIQAGEQASAWIKDFAKNTPLQLDQVTETFVRLKNFGLDPMDGAMQAIIDQSEKL
GGGYERVQGISLALGQAWAKQKLQGEEILQLIERGVPVWQLLENVTGKNTAELQRL
SSAGELGRDTIKQLIDEIGRSAEGSAAKGMSTLSGLVSNARDNFDQFFNLVATSGALD
WLKNQLDSLNKTMAEMAASGELQELAKNISDGIVATAEAVKSLVTTIYEWRGAITA
VGAVWATLKVGSFLADLSKGTLDAIRNLTVLVTTKKGVEIANGKLASSFGPLIGAIRG
GIGAVSDWMKGLSGVGGLLAKGGIFAGIAYGVYEIGRLAKAWLDLREAERALNESR
GEASITNSMVNEELAAINDQLNSNYTSLKEVIAAEEAGQIVREQSTGIWRRNTEEIGR
NTDYLLGHGYALTDSITAIEEAYKSLGLQSTKSLEEAAEASRKAYEVIASGQEPIEQQR
AAFLKYADAANKAAQATGESISDSIKATAANLGLTDSLDKLTGANSKSQVAATEQSK

AMSGASAELAKTKSAIDDYRKTLDSTTASSEEKALAAQQLADAEASLTEQTRRLNEI
KEVEAATYTKLQAKLAEYTQQMQALDELYKADGISAQEYIAQRERYAEVVGIIQRM
LAGLGDGEQKVKEDTDSANLSLAEQQQRLDDLAESSGTATRYISLLANAQQALKTEF
NLTDQTTEDLNKRLNELNGFIVQNNRVTNIWWRELAQASNEAFEREKLIIRETMAMR
GYIQQLGSASLSMAELAQVTKAVDRGFTSLGDNDMAVLRQAITDAENRLLSFRDELE
GTVSSLQDELDRLNDNQAAIEKRAYEQQTAELRAKLAAAQASGDAASIAAAKEALK
LADQIYKTKQAQYAEELKANSKTTNTSSPSSAANNVTPLVRQTTPTSNTVLPTTTASS
TRTVRLVLELQGQSYNADMSISAADQLLAQIERARSTSL*

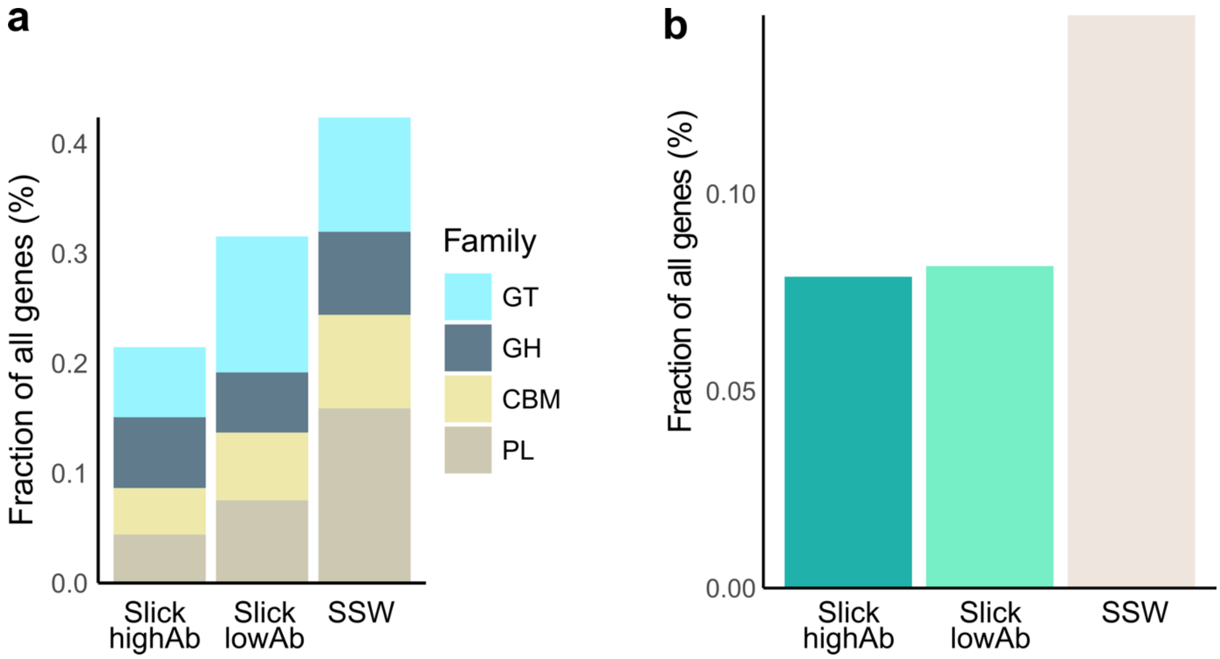


***Fig. S6:*** *Fraction of CAZyme* ***(a)*** *and surfactant-encoding genes in MAG types* ***(b)****, displayed as percent of all encoded genes.*

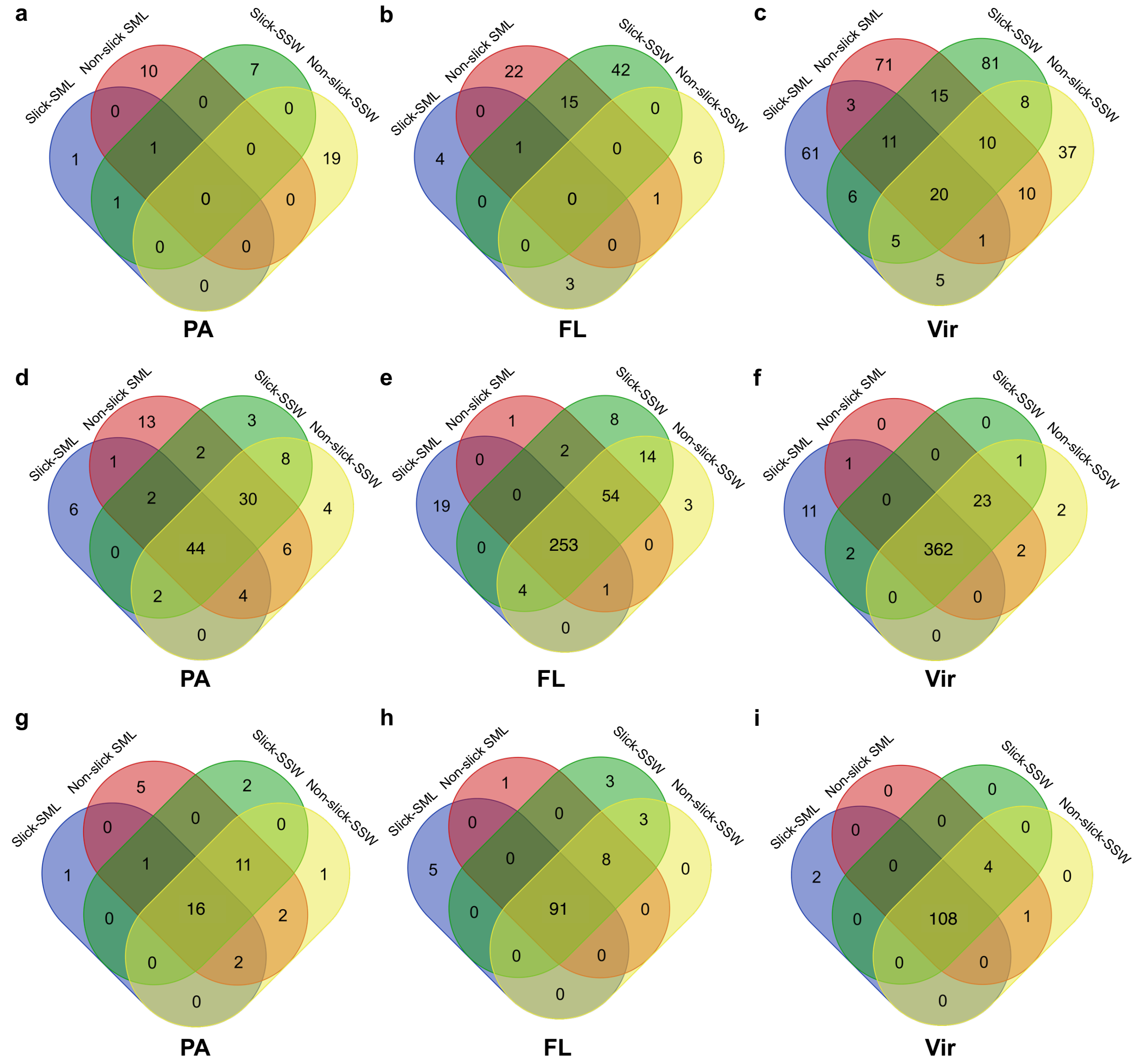


***Fig. S7:*** *Shared vOTUs in viral (<0.2 µm), free-living (FL: 0.2 – 5 µm), particle-associated fraction (PA: >5 µm) for the four sample types. Venn diagrams are based on origin of assembled virus* ***(a-c)****, presence of vOTUs based on read-mapping* ***(d-f)****, and presence of viral clusters (VCs) based on read-mapping* ***(g-i)****. If counts do not add up to 428 (= total number of viral populations detected in this study), this is because not all viruses are found in each fraction. Venn diagrams were constructed using Ugent webtool: [https://bioinformatics.psb.ugent.be/webtools/Venn/](https://bioinformatics.psb.ugent.be/webtools/Venn/). SML = sea-surface microlayer, SSW = subsurface water.*

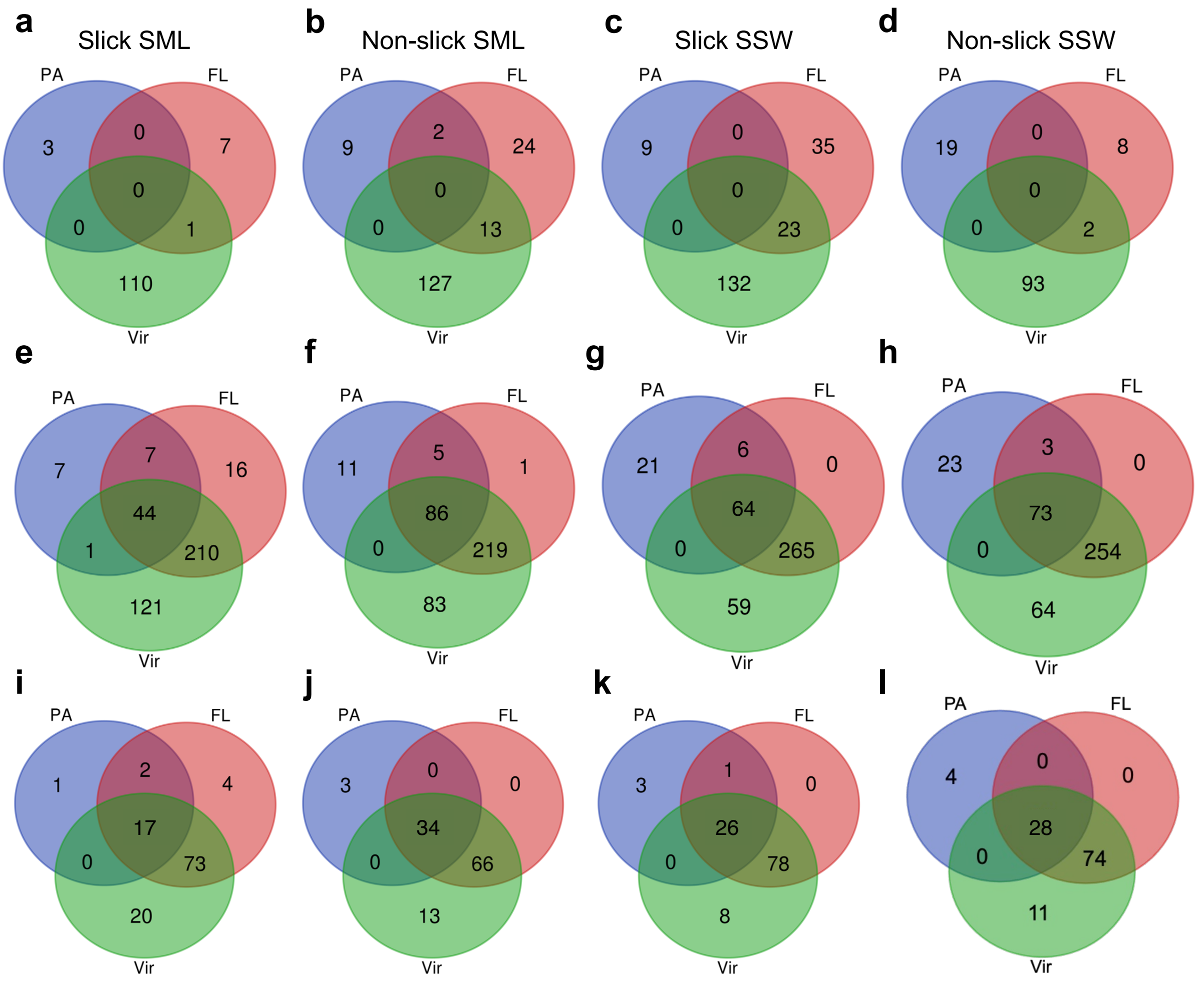


***Fig. S8:*** *Shared vOTUs in slick SML, non-slick SML, slick SSW, and non-slick SSW for the different pore size fractions. Venn diagrams are based on origin of assembled virus* ***(a-d)****, presence of vOTUs based on read-mapping* ***(e-h)****, and presence of viral clusters (VCs) based on read-mapping* ***(i-l)****. If counts do not add up to 428 (= total number of vOTUs detected in this study), this is because not all viruses are found in each fraction. Venn diagrams were constructed using Ugent webtool: [https://bioinformatics.psb.ugent.be/webtools/Venn/](https://bioinformatics.psb.ugent.be/webtools/Venn/). FL = free-living (0.2 – 5 µm), PA = particle-associated fraction (> 5 µm), SML = sea-surface microlayer, SSW = subsurface water.*

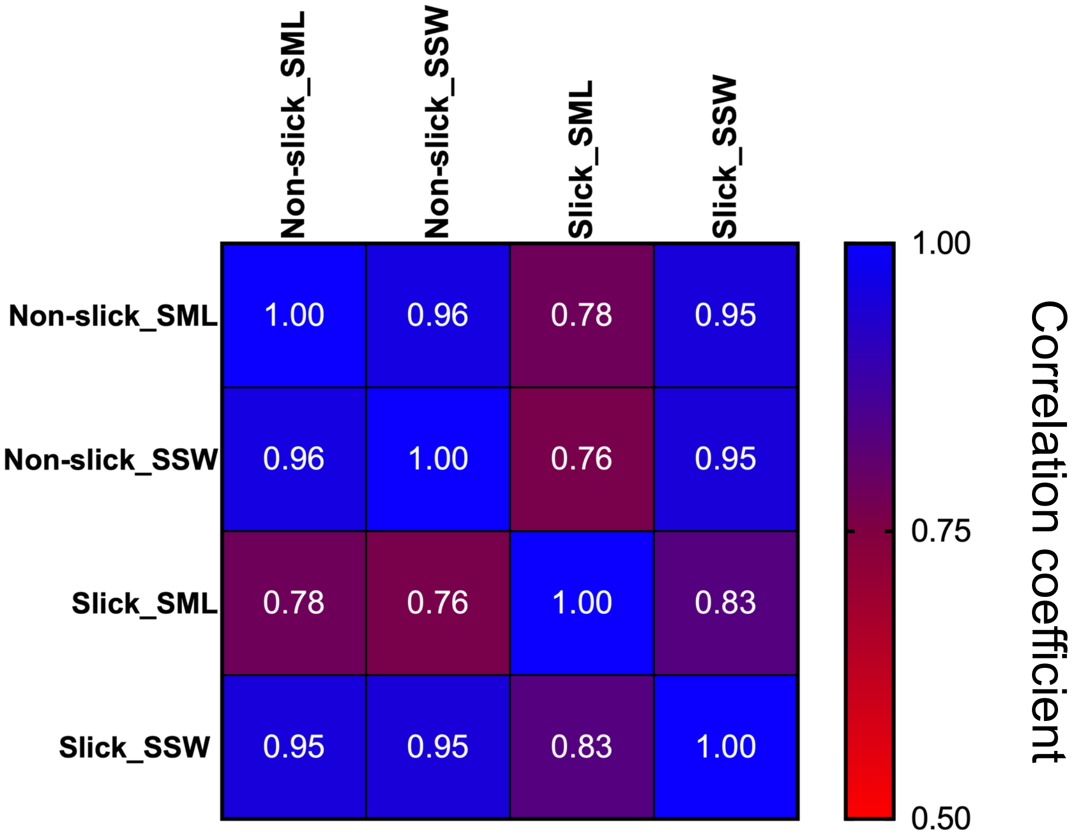


***Fig. S9:*** *Correlation matrix based on read-normalized coverage for 428 viral operational taxonomic units (vOTUs) for the viral fraction (< 0.2 µm) of the samples. Shown is Spearman's R as correlation coefficient. Correlation was performed in GraphPad Prism version 9. SML = sea-surface microlayer, SSW = subsurface water.*

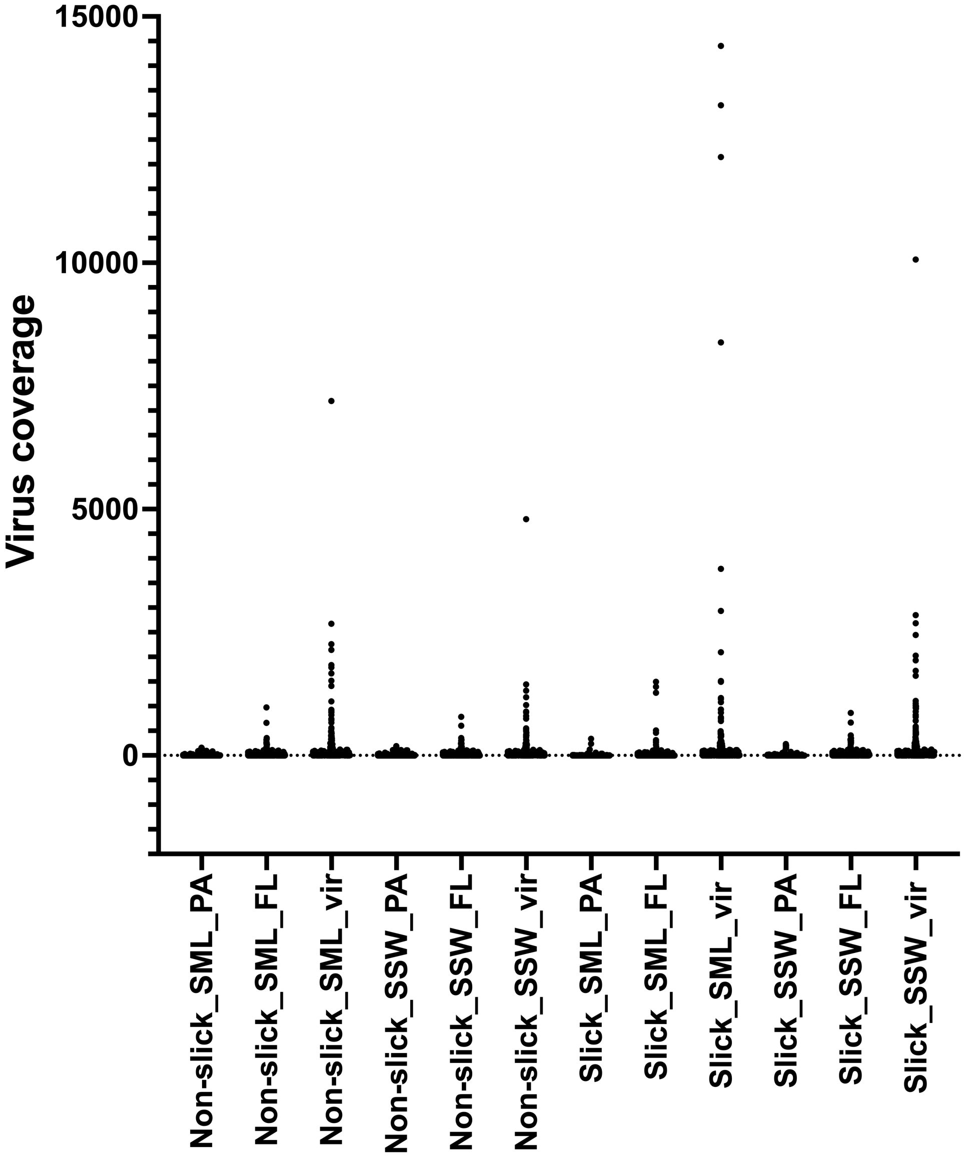


***Fig. S10:*** *Virus coverage normalized to read depth across different samples and filtered fractions with vir (< 0.2 µm), free-living bacterial fraction (FL: 0.2 – 5 µm), particle-associated fraction (PA: >5 µm), SML=sea-surface microlayer, SSW=subsurface water*

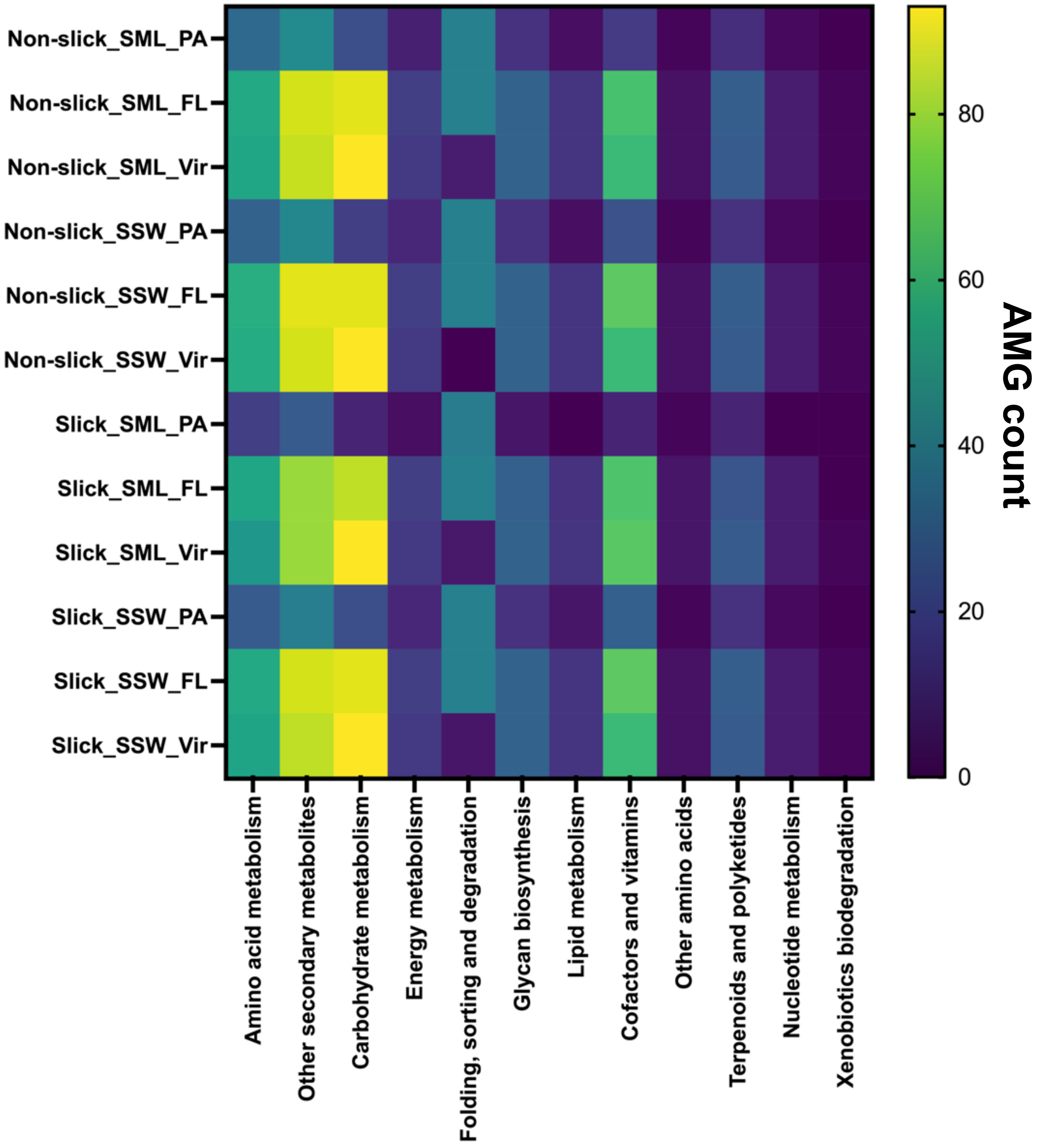


***Fig. S11:*** *Number of viral auxiliary metabolic genes towards a certain metabolic pathway without consideration of viral coverage. FL=free-living fraction (5-0.2 µm pore size filtered), PA=particle-associated fraction (> 5 µm filtered), SML=sea-surface microlayer, SSW=subsurface water (~ 70 cm depth)*

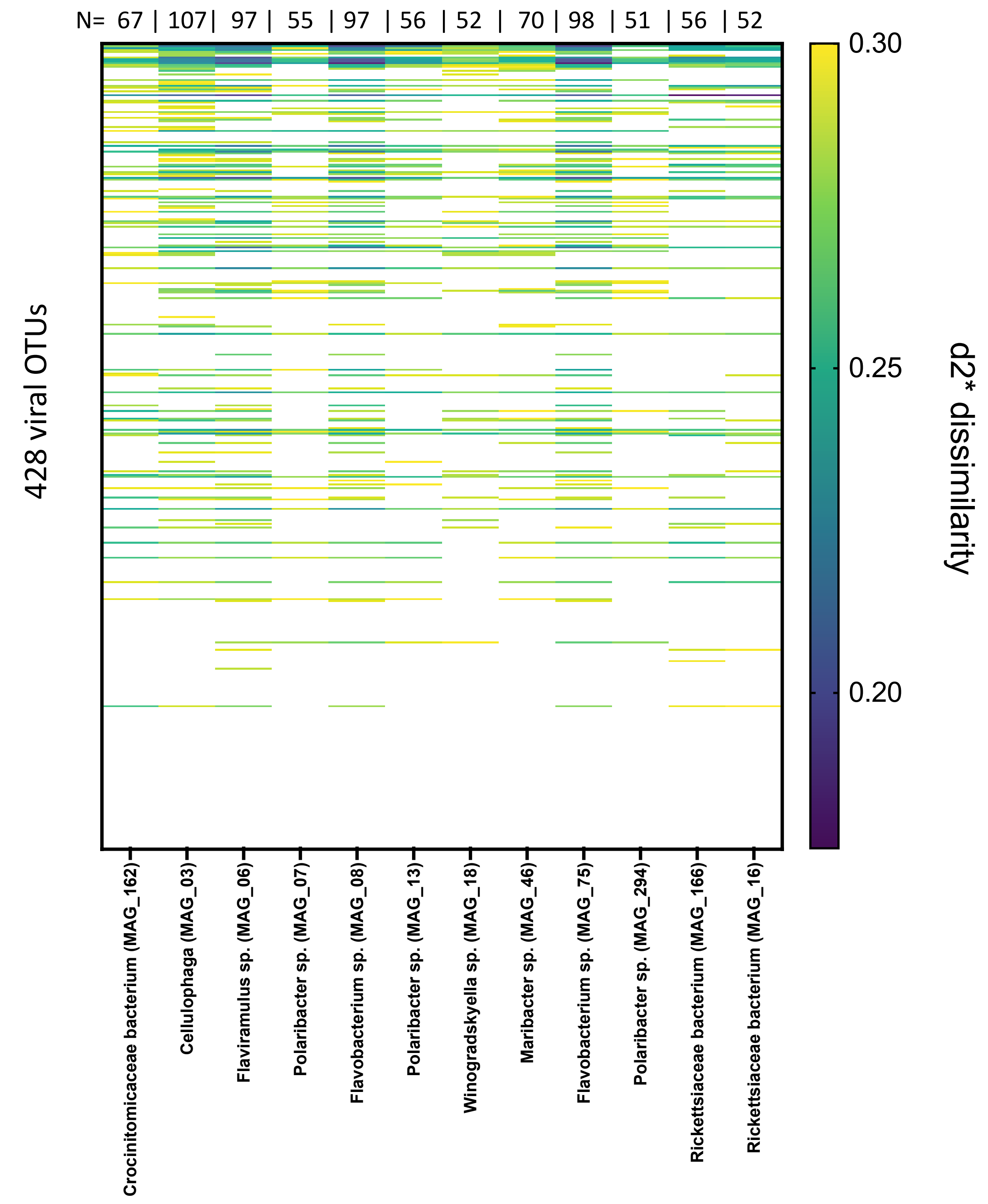


***Fig. S12:*** *Virus-host interactions based on k-mer frequency patterns. Shown are the top 12 host with most k-mer matches to viruses below the d2* threshold of 0.3. Host belonging to orders Flavobacteriales (class Bacteroidia) and Rickettsiales (class Alphaproteobacteria) had most matches with recovered vOTUs. N gives the number of virus-host matches for a metagenome-assembled genome (MAG).*

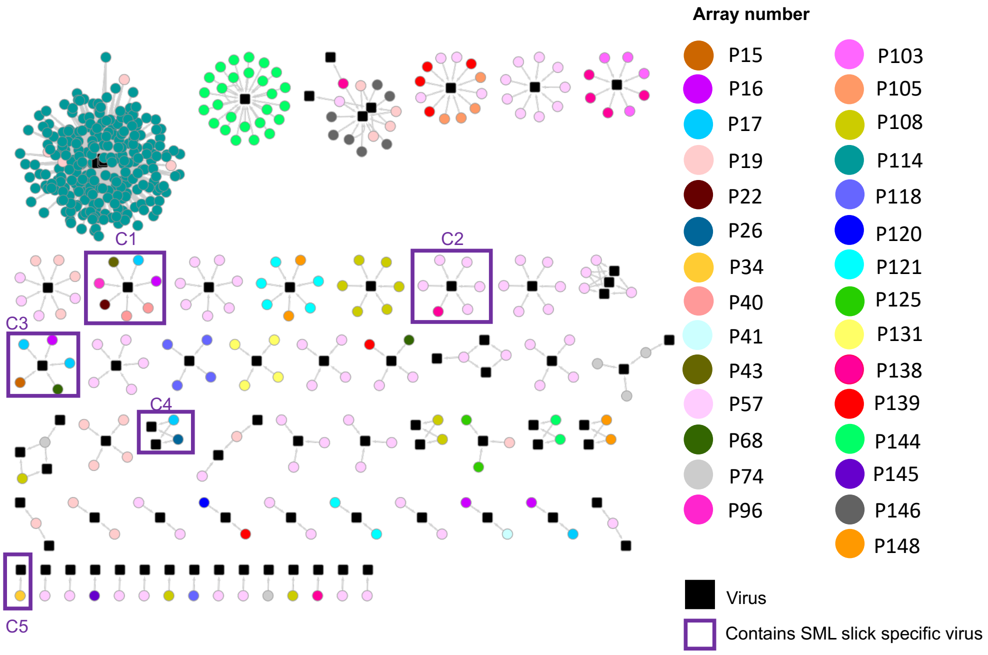


***Fig. S13:*** *Network of CRISPR-spacer to protospacer matches (100% similarity), colored according to CRISPR array the spacer stems from. Purple frames and letters C1- C5 indicate interaction cluster with involvement of viruses only detected in slick SML based on read mapping. Information on spacers matching vOTUs in C1-C5 is given in Table S10.*
